# Controlled release of microorganisms from engineered living materials

**DOI:** 10.1101/2024.09.25.615042

**Authors:** Manivannan Sivaperuman Kalairaj, Iris George, Sasha M. George, Sofía E. Farfán, Yoo Jin Lee, Laura K. Rivera-Tarazona, Suitu Wang, Mustafa K. Abdelrahman, Seelay Tasmim, Asaf Dana, Philippe E. Zimmern, Sargurunathan Subashchandrabose, Taylor H. Ware

## Abstract

Probiotics offer therapeutic benefits by modulating the local microbiome, the host immune response, and the proliferation of pathogens. Probiotics have the potential to treat complex diseases, but their persistence or colonization is required at the target site for effective treatment. Although probiotic persistence can be achieved by repeated delivery, no biomaterial that releases clinically relevant doses of metabolically active probiotics in a sustained manner has been previously described. Here, we encapsulate stiff probiotic microorganisms within relatively less stiff hydrogels and show a generic mechanism where these microorganisms proliferate and induce hydrogel fracture, resulting in microbial release. Importantly, this fracture-based mechanism leads to microorganism release with zero-order release kinetics. Using this mechanism, small (∼1 μL) engineered living materials (ELMs) release >10^8^ colony-forming-units (CFUs) of *E. coli* in 2 h. This release is sustained for at least 10 days. Cell release can be varied by more than three orders of magnitude by varying initial cell loading and modulating the mechanical properties of encapsulating matrix. As the governing mechanism of microbial release is entirely mechanical, we demonstrate controlled release of model Gram-negative, Gram-positive, and fungal probiotics from multiple hydrogel matrices.

**SIGNIFICANCE:** Probiotics offer therapeutic benefits and have the potential to treat complex diseases, but their persistence at the target site is often required for effective treatment. Although probiotic persistence can be achieved by repeated delivery, no biomaterial that releases metabolically active probiotics in a sustained manner has been developed yet. This work demonstrates a generic mechanism where stiff probiotics encapsulated within relatively less stiff hydrogels proliferate and induce hydrogel fracture. This allows a zero-order release of probiotics which can be easily controlled by adjusting the properties of the encapsulating matrices. This generic mechanism is applicable for a wide range of probiotics with different synthetic matrices and has the potential to be used in the treatment of a broad range of diseases.

## INTRODUCTION

Controlled release systems of therapeutic agents aim to deliver a predefined dose at a target site over a sustained period of time^1–5^. Such systems decrease side effects and the number of administrations required for treatment compared to systemic delivery^2^. It is often desirable, but difficult, to generate time-invariant (zero-order) release profiles with the delivered dosage in the therapeutic range^1,2,6,7^. Although fundamental relationships that govern the release kinetics of small molecules and macromolecules from synthetic materials have been developed^1,2^, the governing mechanisms that control the release of microorganisms in a controlled and sustained manner have not been established.

Probiotics are live microorganisms^8,9^ that, when dosed appropriately, offer many health benefits such as modulating the local microbiome^9–13^, the host immune response^14,15^, and the proliferation of pathogens^9,16^. Probiotics have been proposed to provide therapeutic benefits for constipation^17^, diabetes mellitus^18^, *Helicobacter pylori* infections^11^, inflammatory bowel disease^19^, irritable bowel syndrome^19^, *Clostridium difficile* infections^15^, urinary tract infections^10^, and vaginal infections^14^. Although probiotics aid in the treatment of complex diseases, several clinical studies show conflicting results^20,21^, which could be due to the probiotic dosage, mode/timings of administration, strain used, or inclusion of prebiotics^22^. A key challenge to the clinical utility of probiotics is the need to administer probiotics in a metabolically active state at clinically relevant dosage^8^.

Various encapsulation technologies have been developed to increase the survival of probiotics and shield them from harsh environments^3,5,23,24^. For example, in orally delivered probiotics, encapsulation protects the ingested cells from the harsh environment of the stomach^25–27^. However, even if cells are successfully delivered, the persistence of probiotics, an important requirement to achieve beneficial effects^25,28^, is often not observed in the GI tract or other niches^29–32^. To circumvent this drawback, probiotic persistence can be achieved by promoting adhesion between the probiotic and mucus membranes^25^ by either engineering the microbe to enhance its adherence^32^ or coating it with mucoadhesins^33^. For probiotics that lack adhesiveness, repeated administration of a prescribed dose of viable probiotics for a long period can promote colonization^34^. While repeated administration is feasible for accessing some mucosal sites such as the gut and lower reproductive tract, it is a barrier for effective deployment of probiotic-based therapies in hard-to-access sites, such as the urinary bladder^35^.

Currently, controlled release of probiotics is achieved through the degradation of the encapsulating matrices^3–5,8^ or transport of probiotics through porous materials^36^. Although these approaches mirror the mechanisms used for the controlled release of small molecules^2^, they fail to release probiotics in a sustained manner^3–5,8,36^. This is because unlike drug/therapeutic delivery systems where several days’ worth of drug/therapeutics can be loaded and released at the site in a sustained manner^2^, probiotic transport through a solid material is limited due to the large size of cells in comparison to drugs/therapeutics^2,37^, making it difficult to load and deliver many cells. Hence, to release high doses of probiotics in a sustained manner, we need a mechanism that allows local replication of probiotics at the target site, like an in situ probiotic factory^38^.

Engineered Living Materials (ELMs) are composites constructed by embedding living microorganisms into organic or inorganic matrices^13,39–42^. ELMs are endowed with complex emergent functionality from the interplay between their living and non-living components^39,40,43^. The non-living component of ELMs maintains the viability of microorganisms by facilitating the diffusion of water, nutrients, gases, and biomolecules^39^. Notably, the properties of the non-living matrix help modulate the interactions of the microbes with the surrounding environment^41,44^. In many ELMs, microorganism escape is observed^40,43,45,46^, and biocontainment platforms have been developed to prevent that escape^47^. In the ELM field, microbial escape is typically considered a drawback as unwanted release of genetically modified microorganisms and unplanned growth in surroundings could be a source of future regulatory concerns^48^. Whereas, with common probiotics found in the population^44^ or used in food production^49^, microbial release is not a concern. We have previously demonstrated that using ELMs, probiotics can be released from non-porous and non-degrading synthetic polymers^44^. However, the mechanism of microbial escape is poorly understood, and controlled and sustained release of microorganisms has not been demonstrated. Precise control of microorganism escape could be exploited to realize technological or clinical utility.

Herein, we elucidate a generic mechanism in which stiff microorganisms proliferating within relatively low elastic modulus hydrogels induce fracture in those hydrogels causing microbial release. Surprisingly, this mechanism yields sustained release of microorganisms with a zero-order release kinetics. The rate of microbial release is modulated by varying the initial microorganism loading, mechanical properties of the encapsulating hydrogels, and shape and size of the ELMs. Furthermore, since this mechanism is entirely mechanically driven, this mechanism can be extended to several probiotics and synthetic polymers. Hence, we demonstrate the sustained release of a Gram-negative probiotic bacterium (*E. coli* ABU 83972), a Gram-positive probiotic bacterium (*L. paracasei*), and a probiotic fungus (*S. cerevisiae*) from two types of acrylic hydrogels.

## RESULTS AND DISCUSSION

### Microorganisms can be released from ELMs in a sustained manner

To make a probiotic factory and delivery mechanism, we synthesized ELMs with an encapsulated living probiotic, such as *E. coli* (ABU 83972), embedded within acrylic hydrogels. The hydrogels were prepared by free radical polymerization of 2-hydroxyethylacrylate (HEA) (monomer) and *n,n′*-methylenebisacrylamide (BIS) (crosslinker). When these ELMs are incubated in growth media, nutrients diffuse through the hydrogel matrices^39^, allowing bacteria to proliferate within the hydrogel matrices, resulting in colony formation. It is known that colonies exert mechanical forces on the surrounding hydrogel during expansion^41,50^. We hypothesized that upon reaching a certain volume, the colony induces fracture in the hydrogels causing bacterial release (**Fig. 1a, b**). Incubating identical ELMs in non-growth media such as saline does not allow cell growth, hence, no release is observed (**Fig. 1c**). Moving ELMs that were first grown for 5 days into saline dramatically reduces cell release. These data collectively confirm the necessity of microbial proliferation to induce fracture and release microorganisms. Surprisingly, this mechanism leads to sustained release of *E. coli* with a zero-order release kinetics (**Fig. 1c**). ELMs loaded with 1 × 10^4^ cells per μL of ELM are release cells after 1 day of growth and release 4.03 ± 0.18 × 10^7^ colony-forming-units (CFUs) per μL of ELM in 2 h (**Fig. 1c**, **Supplementary Fig. 1**). Moreover, once the ELMs reach steady state release, the number of cells present in the ELMs does not change significantly during the 2 h of cell release (**Fig. 1d**, **Supplementary Fig. 2**). As such, the cells present in the ELMs remain within ELMs to facilitate sustained release.

**Fig. 1.**
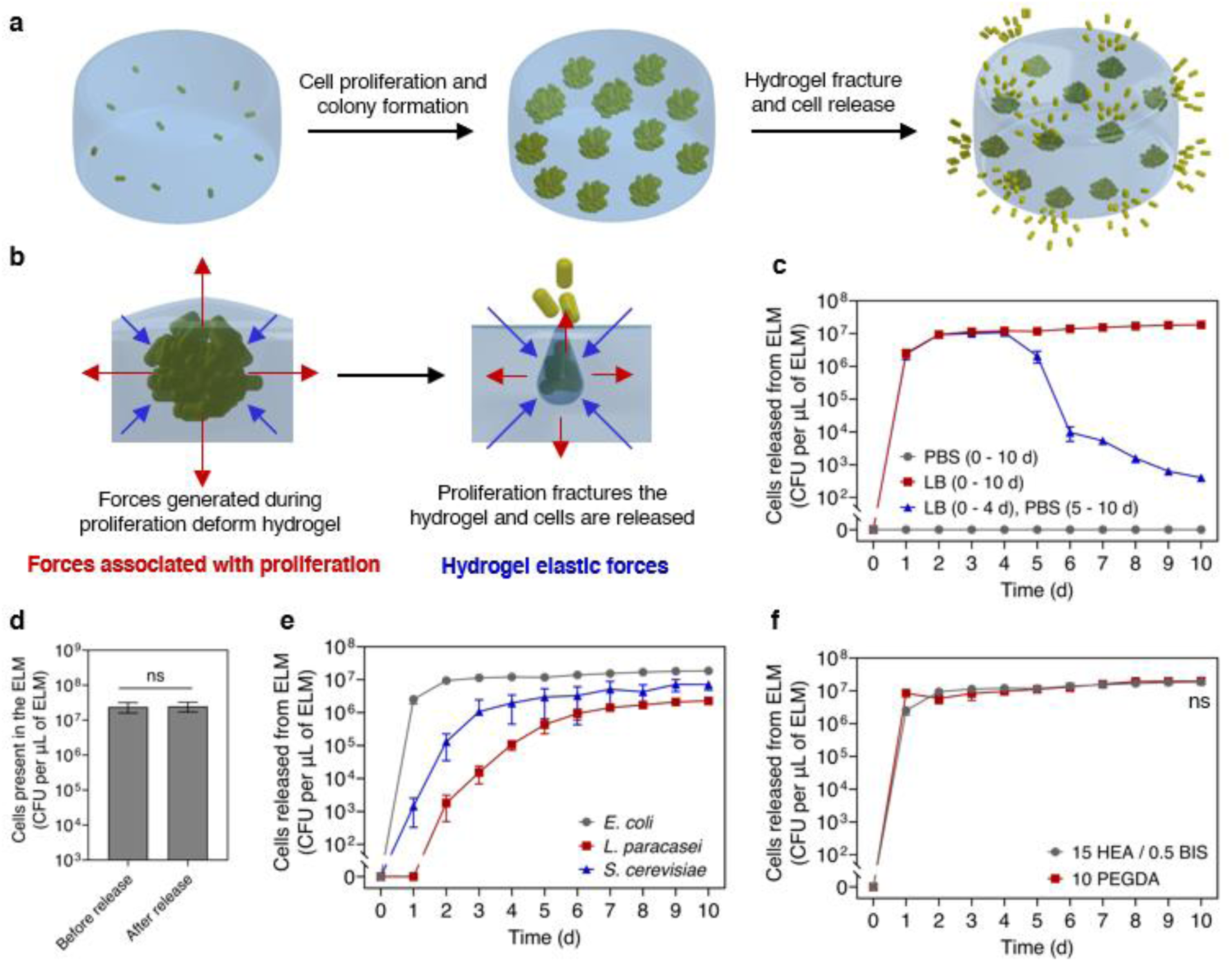
Sustained release of microorganisms from ELMs. **(a)** Schematic illustrating the sustained release of microorganisms from ELMs. Viable microorganisms encapsulated within the ELMs can proliferate, form colonies, and can also be released. **(b)** Schematic illustrating the interplay between microbial proliferation forces and hydrogel elastic forces that induce fracture and cell release. **(c)** Cell release as a function of time when ELMs are incubated in LB media (nutrient media) and saline (PBS). **(d)** The number of cells present within the ELMs before and after 2 h of cell release (5-day grown ELMs). ELMs in panel **c** and **d** were prepared with a cell loading of 1 × 10^4^ *E. coli* ABU 83972 cells per μL of ELM and a hydrogel formulation of 15 HEA / 0.5 BIS (wt%). **(e)** Cell release as a function of time from ELMs loaded with different microorganisms (*E. coli* (gray), *L. paracasei* (red), and *S. cerevisiae* (blue)). All ELMs in panel **d** were prepared with a hydrogel formulation of 15 HEA / 0.5 BIS. *E. coli*-loaded ELMs were prepared with 1 × 10^4^ *E. coli* cells per μL of ELM, *L. paracasei*-loaded ELMs were prepared with 1 × 10^3^ *L. paracasei* cells per μL of ELM, *S. cerevisiae*-loaded ELMs were prepared with 1 × 10^3^ *S. cerevisiae* cells per μL of ELM. All data in panel **c–f** is presented as mean ± standard deviations (*n* = 3). Statistical analysis was performed by a two-tailed Student’s *t*-test. Not significant (ns) for *P* > 0.05.

The driving mechanism for cell release is mechanical, hence, this mechanism can be extended to several probiotics and synthetic polymers. Using the same hydrogel formulations, sustained release of *L. paracasei* and *S. cerevisiae* is observed (**Fig. 1e**). Zero-order release of *E. coli* from ELMs with a different hydrogel matrix made from crosslinking poly(ethylene glycol) diacrylate (**Fig. 1f**). When the stiffness of HEA / BIS ELMs (63.72 ± 1.44 kPa) and PEGDA ELMs (62.36 ± 2.67 kPa) are similar, they show no significant difference in *E. coli* release (*P* > 0.05) (**Fig. 1f**).

### Microorganism growth within ELMs results in the release of microorganisms

Colony expansion induces fracture of encapsulating matrices, resulting in cell release (**Fig. 2a**). To investigate this proposed mechanism, we synthesized ELMs loaded with one colony-forming-unit (*E. coli*). As the relatively stiff *E. coli* (with an elastic modulus > 1 MPa)^51^ multiply within the lower elastic modulus hydrogel matrix (7.52 ± 0.66 kPa), a colony is formed (day 1 to day 5, **Fig. 2b**). Further proliferation increases the colony volume until it induces an observable fracture, resulting in cell release (day 6, **Fig. 2b**). In this case, the shape of the fracture can be seen by the plane of fluorescent cells that remain within the fracture. This plane terminates at the ELM surface. For all ELMs, cell release occurs only after hydrogel fracture (**Fig. 2c**), confirming the necessity of fracture to achieve cell release. As is typical for both biological growth processes and failure processes, the exact time to fracture is somewhat stochastic. The fracture emanating from a growing colony many be similar to cavitation induced failure in polymer networks^52^. As such, fracture likely depends on the exact distance of the growing colony from a free surface.

**Fig. 2.**
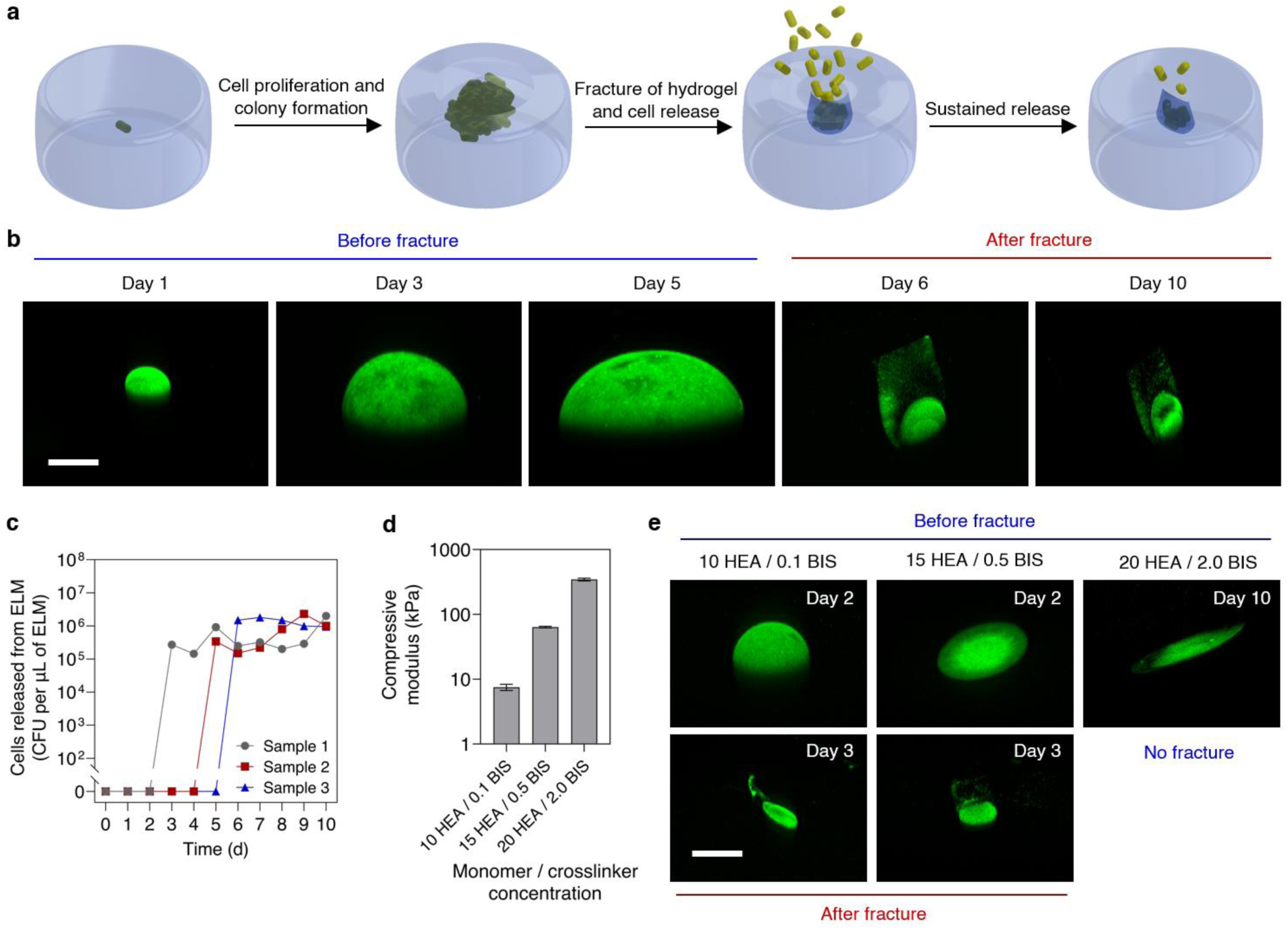
Mechanism for sustained release of microorganisms from ELMs. **(a)** Schematic illustrating the driving mechanism of cell release from ELMs. A single cell encapsulated within an ELM proliferates and forms a colony, which expands until it induces fracture and releases cells. **(b)** Confocal microscopy z-stack images showing colony enlargement and fracture in ELMs. Scale bar, 200 μm. The green fluorescence represents bacteria / bacterial colony and black area represents the encapsulating hydrogel matrix, and image day 1 represents 24 h of incubation. **(c)** Cell release from ELMs as a function of time. ELMs in panels **b** and **c** were 1 mm thick, loaded with single *E. coli*, and prepared with a hydrogel formulation of 10 HEA / 0.1 BIS. **(d)** Compressive modulus of hydrogels with different monomer / crosslinker (HEA / BIS) concentrations. **(e)** Microscopy z-stack images showing colony growth and fracture in ELMs with different stiffnesses (10 HEA / 0.1 BIS, 15 HEA / 0.5 BIS, and 20 HEA / 2.0 BIS). Scale bar, 200 μm. ELMs in panel **e** were 0.5 mm thick and loaded with a single *E. coli*. Data in panel **d** is presented as mean ± standard deviations (*n* = 3).

Although the growth of cells within ELMs is necessary for inducing fracture and releasing cells, cell growth does not always result in fracture and cell release. The ability of the colony to induce fracture depends on the mechanical properties of the encapsulating matrices. We varied the stiffnesses of the hydrogels by varying their monomer (HEA) and crosslinker (BIS) ratios and observed the possibility of ELM fracture and cell release. Increasing the monomer/crosslinker concentrations from 10 HEA / 0.1 BIS to 15 HEA / 0.5 BIS and 20 HEA / 2.0 BIS increases the compression modulus of the hydrogel from 7.52 ± 0.66 kPa to 63.72 ± 1.44 kPa and 345.16 ± 14.29 kPa, respectively (**Fig. 2d**, **Supplementary Fig. 3**). At all stiffnesses, bacterial proliferation leads to colony formation and expansion within ELMs (**Fig. 2e**). However, the stiffness of the hydrogels dictates the size and morphology of the colonies, as has been seen in other ELMs^41,50^. The volume of the colony in low-stiffness ELMs is significantly larger than the medium-stiffness ELMs and high-stiffness ELMs (**Supplementary Fig. 4a**). In low-stiffness ELMs the colony morphology is spherical while in high-stiffness ELMs the colony morphology is plate-like. Likely, this change in morphology minimizes the elastic strain energy of the growing inclusion^53^. The colony in the high-stiffness hydrogels is unable to cause fracture during the entire period. Therefore no cell release is observed (**Supplementary Fig. 4b**). This suggests that the high stiffness of hydrogel restricts the ability of the colony to reach a volume that can induce hydrogel fracture. This result also demonstrates that cell release can be tuned by varying the properties of the encapsulating matrices.

The volume of the encapsulating matrix plays a major role in allowing fracture and subsequent cell release. Colonies encapsulated within a smaller volume of low stiffness hydrogel have a better ability to induce hydrogel fracture compared to a larger volume of the same encapsulating matrix. We synthesized single-cell ELMs with different thicknesses (0.5, 1, and 2 mm) and observed if the colony expansion induced hydrogel fracture. At all thicknesses, bacterial growth leads to colony formation and expansion within ELMs (**Supplementary Fig. 5a**). However, only 0.5 mm and 1 mm thick ELMs undergo fracture with colony expansion. Moreover, 1 mm thick ELMs require a significantly larger colony to induce fracture compared to 0.5 mm thick ELMs (**Supplementary Fig. 5b**). Hence, 1 mm thick ELMs demonstrate a delayed fracture and delayed cell release. The hydrogel fracture in both 0.5 mm and 1 mm thick ELMs allows cell release in a sustained manner (**Supplementary Fig. 5c**). On the other hand, colonies in 2 mm thick ELMs are unable to induce fracture throughout the entire experiment (10 days). The larger hydrogel is better able to accommodate the strain caused by the growing inclusion (colony) without failure. Critically, due to the absence of fracture, these ELMs do not release cells (**Supplementary Fig. 5c**), highlighting the importance of fracture to achieve cell release. These model ELMs with only a single colony help in understanding the mechanism of cell release from ELMs, but the number of released cells is somewhat difficult to control (**Supplementary Fig. 4b, 5c**).

### Hydrogel stiffness controls microorganism release

The stiffness of the encapsulating hydrogel matrix controls cell proliferation, colony formation, and cell release from ELMs with many embedded cells. To increase the number of cells being released, we first increased the cell loading to 1 × 10^4^ cells per μL of ELM. The presence of a higher number of cells within ELMs increases the number of colonies and, likely, the number of fractures. In turn, cell release is increased as compared to the ELMs with a single colony. We synthesized hydrogels with the same cell loading (1 × 10^4^ cells per μL of ELM) but different stiffnesses by varying their formulations (10 HEA / 0.1 BIS (low stiffness), 15 HEA / 0.5 BIS (medium stiffness), and 20 HEA / 2.0 BIS (high stiffness)) and quantified their cell release. To determine if the monomers and crosslinkers used during the preparation of hydrogels produced any cytotoxic effect, we performed toxicity tests. There is no significant difference in the cell viability when they are exposed to different concentrations of monomer and crosslinker (*P* > 0.05) (**Supplementary Fig. 6**). Like the single-colony ELMs, the colony volume in these ELMs increases with time but decreases with hydrogel stiffnesses (**Fig. 3a).** Low stiffness ELMs have a significantly higher colony volume compared to both medium stiffness (*P* < 0.0001) and high stiffness ELMs (*P* < 0.0001) (**Fig. 3b**). The number of viable cells present in the ELMs also increases with time. The number of cells present at day 10 within low stiffness ELMs (2.39 ± 0.18 × 10^8^ CFUs per μL of ELM) is significantly higher than both medium stiffness (7.45 ± 0.26 × 10^7^ CFUs per μL of ELM, *P* < 0.0001) and high stiffness ELMs (4.03 ± 0.18 × 10^7^ CFUs per μL of ELM, *P* < 0.0001) (**Fig. 3c**). For all stiffnesses, the ELMs demonstrate release after 1 day, and within 3 days, the release reached a steady state (a zero-order release kinetics, **Fig. 3d**). This steady state release remained for the duration of the experiment, which we end arbitrarily after 10 days. The increase in the cell release observed during the first three days could be attributed to new fractures occurring during that time. Whereas the steady state observed after 3 days could indicate the lack of new fractures. During the steady state, the number of cells being released is proportional to the number of cells present within the ELMs and is primarily controlled by the doubling time of the microorganism. On day 10, the low-stiffness, medium-stiffness, and high-stiffness ELMs released 2.24 ± 0.06 × 10^8^, 1.85 ± 0.08 × 10^7^, and 3.14 ± 0.16 × 10^6^ CFUs per μL of ELM, respectively (**Fig. 3d**). The low-stiffness ELMs release significantly more cells than both medium-stiffness (*P* < 0.0001) and high-stiffness ELMs (*P* < 0.0001), and the medium-stiffness ELMs release significantly more cells than high-stiffness ELMs (*P* < 0.01). We note that the cells released from the ELMs can proliferate in the surrounding media, however, proliferation in the media does not significantly influence the quantity measured (**see Supplementary text, and Supplementary Fig. 7**). The cell release is driven by the forces associated with cell proliferation, and not influenced by other external forces, such as those arising from shaking (**see Supplementary text, and Supplementary Fig. 8**).

**Fig. 3.**
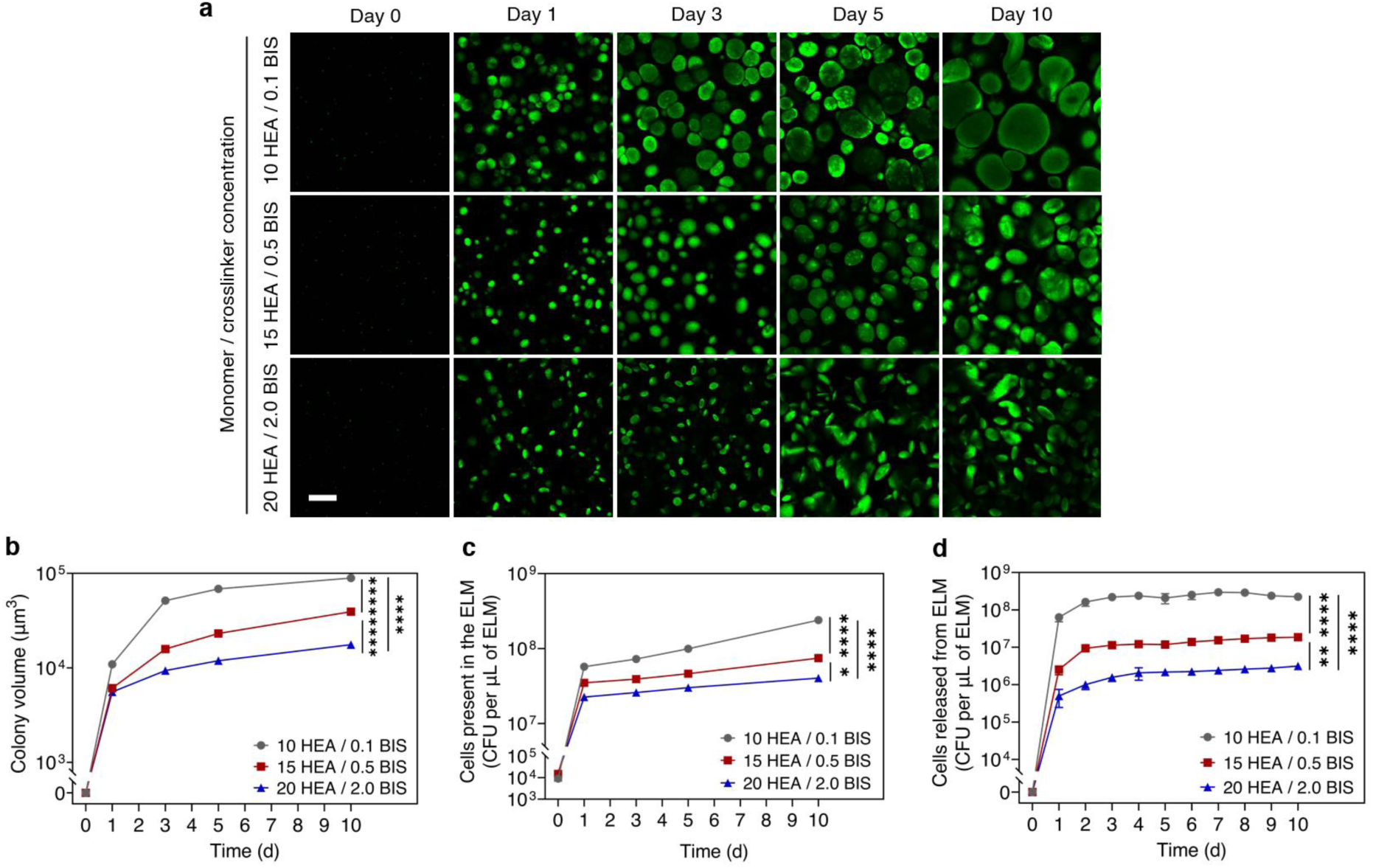
Hydrogel stiffness controls microorganism release. **(a)** Confocal microscopy images showing the differences in colony morphologies as the hydrogel stiffness is varied. Scale bar, 100 μm. **(b)** Colony volumes as a function of time for ELMs with different stiffnesses. **(c)** The number of cells present within ELMs as a function of time for ELMs prepared with different hydrogel stiffnesses. **(d)** Cells released from ELMs as a function of time for ELMs prepared with different hydrogel stiffnesses. Hydrogel stiffnesses were varied by varying the hydrogel formulations (10 HEA / 0.1 BIS (gray), 15 HEA / 0.5 BIS (red), and 20 HEA / 2.0 BIS (blue)). All ELMs were prepared with a cell loading of 1 × 10^4^ cells per μL of ELM. All data in panel **b** is presented as mean ± standard error of means (*n* > 1000) and all data in panel **c–d** is presented as mean ± standard deviations (*n* = 3). Statistical analysis was performed by a one-way ANOVA with post-hoc Tukey’s test, * *P* < 0.05, ** *P* < 0.01, and **** *P* < 0.0001.

The ease of controlling probiotic release by just varying the stiffness of the encapsulating matrix demonstrates the potential of this technology for different types of diseases where the probiotic dose is crucial. These small (∼1 μL) ELMs release >10^8^ CFUs in 2 h, demonstrating the ability to release high doses of probiotics in a short period, which is necessary for therapeutic efficacy^54,55^. Moreover, the release of high doses of probiotics is sustained for days. Since repeated administration of a prescribed dose of viable probiotics for a long period can promote colonization^34^, the ability of this approach to release high doses in a sustained manner for a prolonged time makes ELM a good candidate for achieving probiotic persistence/colonization.

### Microorganism release can be tuned by initial cell loading

Cell release can be varied by controlling the number of cells present in the ELMs. We quantified the cells released from ELMs synthesized with the same stiffness (15 HEA / 0.5 BIS) but different initial cell loadings (10^0^ cells per μL of ELM, 10^4^ cells per μL of ELM, and 10^8^ cells per μL of ELM). Increasing the cell loading increases the number of viable cells (**Supplementary Fig. 10**), which in turn increases the number of colonies present within the ELMs (**Fig. 4a**). For ELMs with 10^8^ cells per μL of ELM (high-cell loading), imaging individual colonies and quantifying colony volumes was not possible due to near confluent growth (**Fig. 4a**). Over 10 days of growth, all ELMs demonstrate an increase in the number of cells present within the ELMs (**Fig. 4b**). Comparing ELMs with different cell loading after 10 days show that high-cell loading ELMs have a significantly higher number of cells (1.6 ± 0.26 × 10^8^ CFUs per μL of ELM) than both medium-cell loading (7.45 ± 0.26 × 10^7^ CFUs per μL of ELM, *P* < 0.01) and low-cell loading ELMs (6.68 ± 2.54 × 10^6^ CFUs per μL of ELM, *P* < 0.001) (**Fig. 4b**). Similarly, medium-cell loading ELMs has significantly more cells than low-cell loading ELMs (*P* < 0.05). For high and medium-cell loading, the ELMs reached a steady state release within 3 days and sustained this release for the remainder of the experiment (**Fig. 4c**). Whereas low-cell loading ELMs demonstrate a delay in reaching a steady state release (6 days). On day 10, the low, medium, and high-cell loading ELMs released 8.93 ± 0.34 × 10^5^, 1.85 ± 0.08 × 10^7^, and 3.34 ± 0.05 × 10^8^ CFUs per μL of ELM, respectively. The high-cell loading ELMs release significantly higher cells than both medium (*P* < 0.0001) and low-cell loading ELMs (*P* < 0.0001), and the medium-cell loading ELMs release significantly more cells than low-cell loading ELMs (*P* < 0.01). These data collectively suggest that cell release can be controlled by varying cells present within the ELMs.

**Fig. 4.**
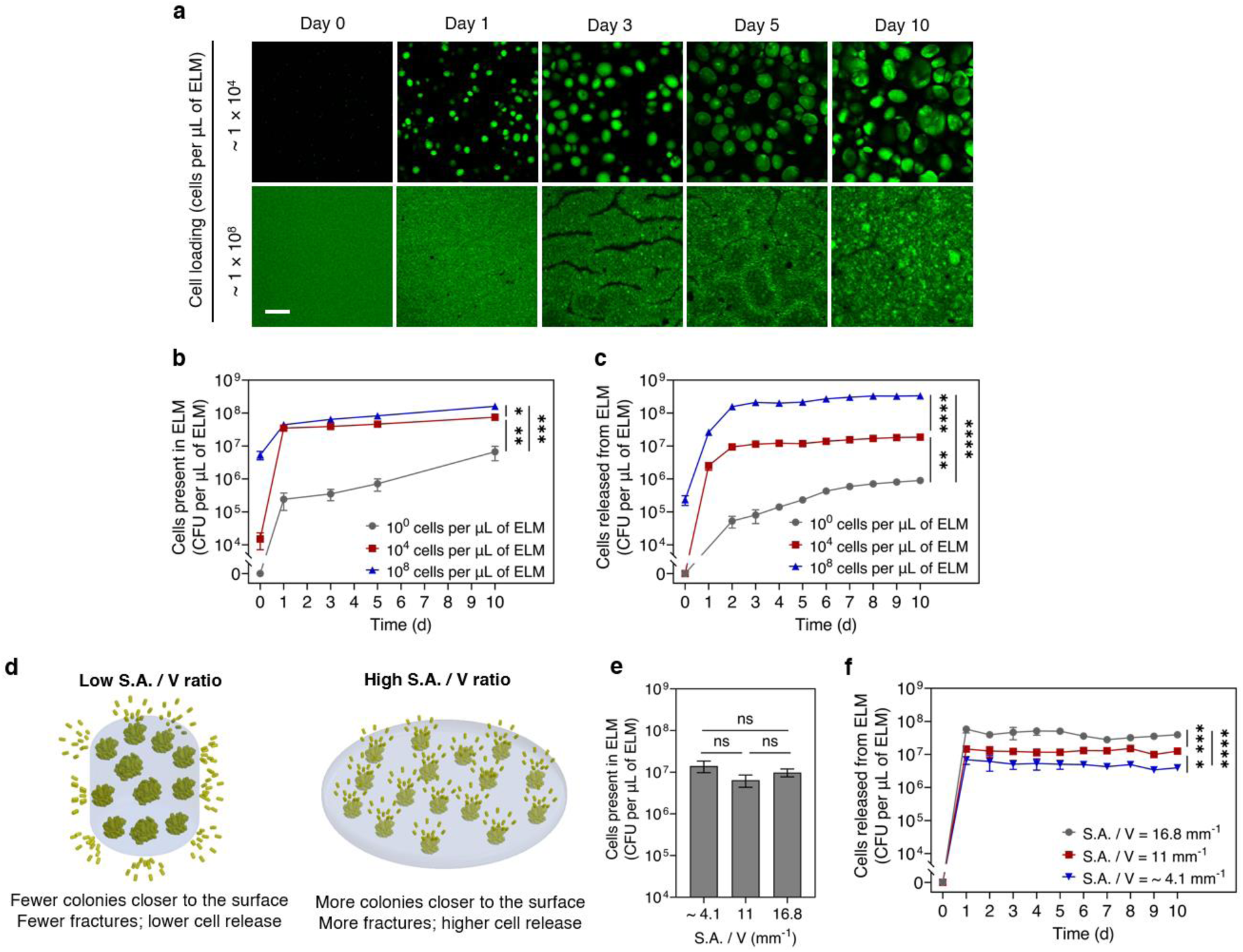
Controlling microorganism release by varying cell loading and shape of ELMs. **(a)** Confocal microscopy images showing cell proliferation and colonies within ELMs of different cell loading. Scale bar, 100 μm. **(b)** The number of cells present within ELMs as a function of time for ELMs prepared with different cell loading. **(c)** Cells released from ELMs as a function of time for ELMs prepared with different cell loading. ELMs in panel **b** and **c** were prepared with 15 HEA / 0.5 BIS formulation and their cell loading was varied from 1 × 10^0^ cells per μL of ELM (gray) to 1 × 10^4^ cells per μL of ELM (red) and 1 × 10^8^ cells per μL of ELM (blue). The initial volume and surface-area-to-volume (S.A./V) ratios of all the ELMs in panel **a–c** were the same. **(d)** Schematic showing influence of surface-area-to-volume (S.A./V) ratio on cell release. ELMs with a high S.A./V ratio have more colonies closer to the surface, which induces more fracture, thereby releasing more cells than ELMs with low S.A./V ratio. **(e)** The number of cells present within ELMs after 10 days of cell release for ELMs prepared with different S.A./V ratios (16.8 mm^−1^, 11 mm^−1^, and ∼4.1 mm^−1^). **(f)** Cell release as a function of time from ELMs with different S.A./V ratios (16.8 mm^−1^ (gray), 11 mm^−1^ (red), and ∼4.1 mm^−1^ (blue)). ELMs in panels **e** and **f** were prepared with a cell loading of 1 × 10^4^ cells per μL of ELM and 15 HEA / 0.5 BIS formulation. All data are presented as mean ± standard deviations (*n* = 3). Statistical analysis was performed by a one-way ANOVA with post-hoc Tukey’s test. * *P* < 0.05, ** *P* < 0.01, *** *P* < 0.001, **** *P* < 0.0001, and not significant (ns) for *P* > 0.05.

### Size and shape of ELMs control microorganism release

Not all colonies within ELMs induce hydrogel fracture and facilitate cell release. We hypothesized that the colonies that are closer to the surface have a higher probability of inducing rupture and therefore releasing cells (**Fig. 4d**). We compared the cell release from ELMs with the same volume (2.48 ± 0.24 mm^3^) but different surface-area-to-volume (S.A./V) ratios (**Fig. 4d**). Varying the S.A./V ratios does not significantly change the number of cells present in ELMs over 10 days of growth (**Fig. 4e, Supplementary Fig. 11**). Moreover, ELMs with different S.A./V ratios all demonstrate zero-order release kinetics for the entire period (**Fig. 4f**). However, increasing the S.A./V ratio from ∼4.1 to 11 and 16.8 mm^−1^ increases the cell release from 3.93 ± 0.66 × 10^6^ CFUs per μL of ELM to 1.25 ± 0.13 × 10^7^ CFUs per μL of ELM and 3.93 ± 0.41 × 10^7^ CFUs per μL of ELM, respectively (**Fig. 4f**). ELMs with an S.A./V ratio of 16.8 mm^−1^ release significantly more cells than ELMs with an S.A./V ratio of 11 mm^−1^ (*P* < 0.001) and ∼4.1 mm^−1^ (*P* < 0.0001). Similarly, ELMs with an S.A./V ratio of 11 mm^−1^ also release significantly more cells than ELMs with an S.A./V ratio of ∼4.1 mm^−1^ (*P* < 0.05). These data suggest that colonies closer to the surface play a significant role in cell release. This phenomenon is somewhat expected due to potentially higher proliferation near the surface and the increased propensity for the growing colony near the surface to induce failure^41,56^.

ELM size also plays a role in controlling cell release. To test the controllability of cell release from ELMs of different sizes, we varied the volume of the ELMs (2.47 and 105.5 mm^3^) without changing the S.A./V ratios (4.08 ± 0.02 mm^−1^). The smaller ELMs (2.47 mm^3^) demonstrate a zero-order release kinetics after 1 day. Whereas the larger ELMs (105.5 mm^3^) demonstrate only reach zero-order release kinetics after 3 days (**Supplementary Fig. 12**). Moreover, as the initial volume of the ELMs increases from 2.47 to 105.5 mm^3^, the cell release from the ELMs reduces significantly from 3.93 ± 0.66 × 10^6^ CFUs per μL of ELM to 6 ± 1.91 × 10^5^ CFUs per μL of ELM (*P* < 0.05). The reason for reduced cell release for larger ELMs could be due to the inherent length scales associated with diffusion and consumption of nutrients, or the volume of media could also be disproportionally small for the volume of ELMs. Hence, this could induce competition between the cells present within the ELMs, resulting in further reduced cell release.

### Other microorganisms can be released via the same mechanism

Since cell release is governed by mechanics and not chemical mechanisms, this mechanism can be extended to a wide range of probiotics. Moreover, the release of those probiotics can also be controlled using the same techniques such as varying the stiffness of the encapsulating matrices and the initial cell loading. However, the rate of proliferation, size, and mechanical properties of the cells are controlled by the genetics of the organism, which leads to changes in the rate of release^9,16,57,58^. We demonstrate the controlled release of probiotics that belong to a different bacterial phylum (*L. paracasei*, a Gram-positive bacterium) and a different kingdom (*S. cerevisiae*, a fungus). All of these organisms have higher stiffness relative to the synthesized hydrogels^37,51,58^. We first synthesized ELMs loaded with *L. paracasei* (1 × 10^3^ cells per μL of ELM) with different stiffnesses by varying their formulations (10 HEA / 0.1 BIS (low stiffness), 15 HEA / 0.5 BIS (medium stiffness), and 20 HEA / 2.0 BIS (high stiffness)) and quantified their cell release (**Fig. 5a**). The low-stiffness ELMs release significantly more cells than both medium-stiffness (*P* < 0.0001) and high-stiffness ELMs (*P* < 0.0001). Next, we quantified the cell release from medium stiffness ELMs (15 HEA / 0.5 BIS) loaded with different cell loadings (10^0^ cells per μL of ELM (low-cell loading), 10^3^ cells per μL of ELM (medium-cell loading), and 10^6^ cells per μL of ELM (high-cell loading)) (**Fig. 5b**). The high-cell loading ELMs release significantly more cells than both medium (*P* < 0.0001) and low-cell loading ELMs (*P* < 0.0001). Similarly, comparing cell release from *S. cerevisiae* loaded ELMs (1 × 10^3^ cells per μL of ELM) with different hydrogel stiffnesses (10 HEA / 0.1 BIS (low stiffness), 15 HEA / 0.5 BIS (medium stiffness), and 20 HEA / 2.0 BIS (high stiffness)) show that low stiffness ELMs release significantly more cells than both medium-stiffness (*P* < 0.0001) and high-stiffness ELMs (*P* < 0.0001) (**Fig. 5c**). Also, comparing cell release from *S. cerevisiae* loaded ELMs (15 HEA / 0.5 BIS (medium stiffness)) with different cell loadings (10^0^ cells per μL of ELM (low-cell loading), 10^3^ cells per μL of ELM (medium-cell loading), and 10^6^ cells per μL of ELM (high-cell loading)) show that high-cell loading ELMs release significantly more cells than both medium (*P* < 0.0001) and low-cell loading ELMs (*P* < 0.0001) (**Fig. 5d**). Compared to *E. coli*-loaded ELMs, a delay in cell release is observed in both *L. paracasei*-loaded ELMs and *S. cerevisiae*-loaded ELMs. Although both *L. paracasei*-loaded ELMs and *S. cerevisiae*-loaded ELMs demonstrate zero-order release kinetics, they require a longer time to reach the steady-state release. The delay in both the start of release and the time taken to take steady-state could be attributed to the slow doubling time of *L. paracasei* and *S. cerevisiae* compared to *E. coli*^9,16,57,58^.

**Fig. 5.**
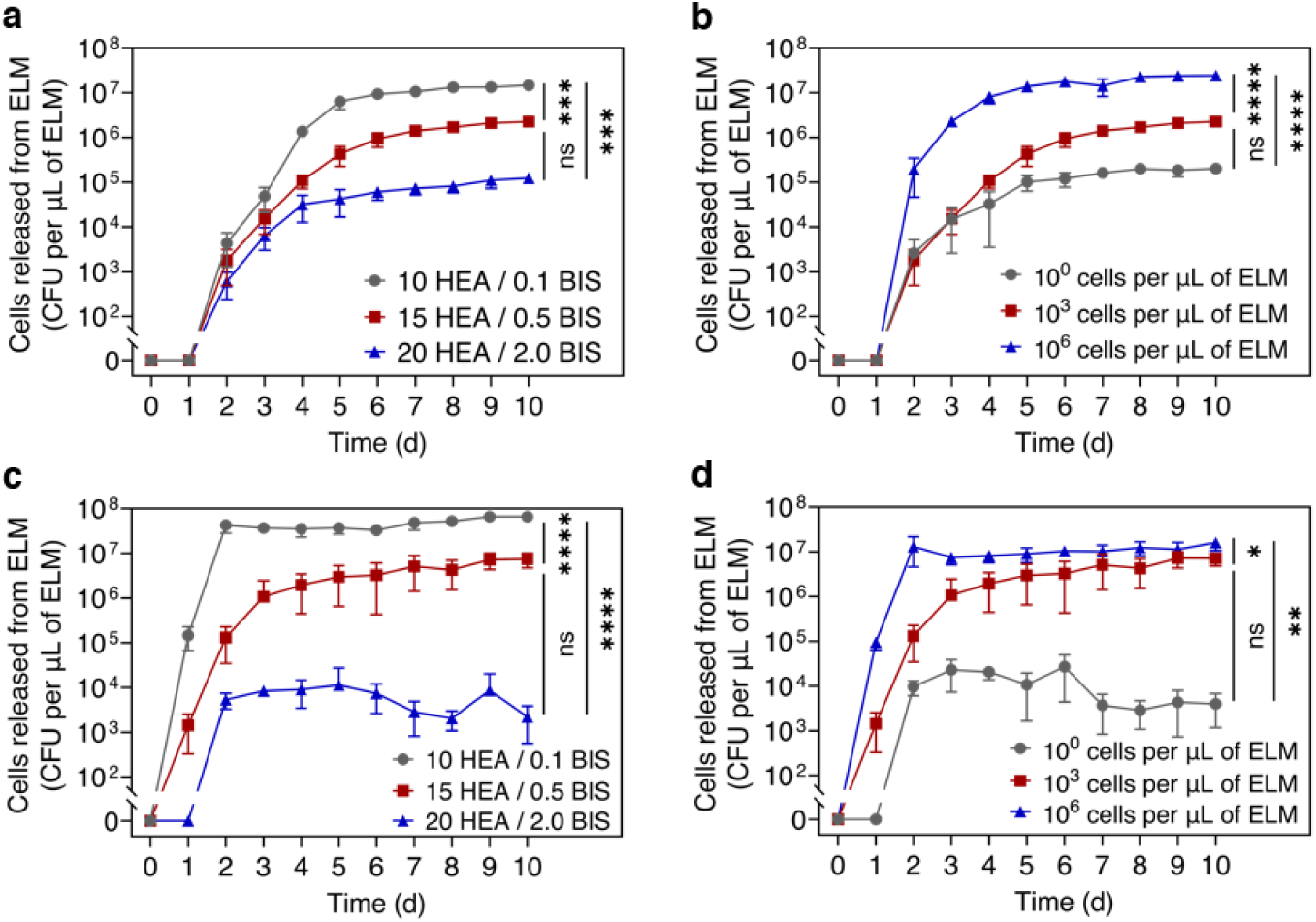
Controlled release of other microorganisms from ELMs. (a, **b)** Controlled release of *L. paracasei* by varying hydrogel stiffnesses and initial cell loading. **(a)** Cell release as a function of time from ELMs with different hydrogel stiffnesses. **(b)** Cell release as a function of time from ELMs with different cell loading. **(c, d)** Controlled release of *S. cerevisiae* by varying hydrogel stiffnesses and initial cell loading. **(c)** Cell release as a function of time from ELMs with different hydrogel stiffnesses. **(d)** Cell release as a function of time from ELMs with different cell loading. All ELMs in panels **a** and **c** were prepared with a cell loading of 1 × 10^3^ cells per μL of ELM, and all ELMs in panels **b** and **d** were prepared with 15 HEA / 0.5 BIS. All data are presented as mean ± standard deviations (*n* = 3). Statistical analysis was performed by a one-way ANOVA with post-hoc Tukey’s test. * *P* < 0.05, ** *P* < 0.01, *** *P* < 0.001, **** *P* < 0.0001, and not significant (ns) for *P* > 0.05.

Using this fracture-based mechanism, different types of probiotics can be released from ELMs. In this paper, we demonstrated controlled release of *E. coli* (ABU 83972), *L. paracasei*, and *S. cerevsiae* from ELMs. ABU 83972 is an *E. coli* strain (Gram-negative bacterium) that offers superior protection against recurrent urinary tract infections (UTIs) compared to the current standard of treatment (antibiotic therapy)^16,59^, representing an alternative approach to antibiotics for preventing UTIs. ABU 83972 outcompetes uropathogenic *E. coli* and other common uropathogens, but do not consistently persist or colonize the bladder^9,16,35^. Hence, a bladder-resident ELM that releases ABU 83972 in a sustained manner could overcome this limitation, thereby conferring a potential alternative for treating UTIs and recurrent UTIs. Several probiotics have persistence or colonization limitations, which can be overcome with this approach. For example, *L. paracasei* is a Gram-positive bacterium that have been used as probiotics for treating UTIs^12^, bacterial vaginosis^60^, and fungal infections^60^. Similarly, *S. boulardii* and *S. cerevisiae* are probiotics that belong to the fungal kingdom and have shown potential in treating diarrhea^61^, colitis^61^, bacterial vaginosis^62^, and candida infections^62^. These probiotics also have persistence limitations^63,64^; hence, using a material that releases these probiotics in a sustained manner could overcome such limitations.

## CONCLUSIONS

We report a simple yet powerful method to release microorganisms in a sustained and controlled manner using ELMs. In these ELMs, the encapsulating hydrogel matrix maintains the viability of the microorganisms and allows microbial proliferation into colonies. Further proliferation expands the colonies and induces fracture, resulting in microbial release. Using single-cell ELMs, we demonstrate that the cell location, along with the stiffness and volume of the encapsulating matrix, dictate hydrogel fracture. We control cell release by varying initial cell loading and encapsulating matrix properties such as stiffness, size, and shape. Since the driving mechanism for this approach is mechanical, this mechanism can be extended to several probiotics and synthetic polymers. Hence, we demonstrate sustained release of *E. coli* (Gram-negative bacterium), *L. paracasei* (Gram-positive bacterium), and *S. cerevisiae* (fungus) for up to 10 days. We also show release of microorganisms from two different classes of acrylic hydrogels. Although probiotics demonstrate great potential in treating complex diseases, lack of persistence or colonization reduces the effectiveness of these therapies. This ELM-based technique may allow for the persistent delivery of a wide variety of probiotics at a target site and aid in colonization, thereby improving treatment efficacy. Furthermore, the simple fabrication technique and release mechanism may enable large scale fabrication and translating this technology to practical applications.

## METHODS

### Materials

2-Hydroxyethylacrylate (HEA), *N,N′*-methylenebisacrylamide (BIS), polyethylene glycol diacrylate (PEGDA, MW = 700 g mol^−1^), and Lithium phenyl-2,4,6-trimethylbenzoylphosphinate (LAP) were purchased from Sigma-Aldrich. ABU 83972 strain of *E. coli* was obtained from Subash lab^16,65^, *L. paracasei* D3–5 strain was obtained from Yadav lab^66^, and *S. cerevisiae* (Active Dry) was obtained from Fleischmann’s Yeast. Powdered forms of Luria-Bertani (LB) Lennox broth, De Man, Rogosa, and Sharpe (MRS) broth, Yeast Peptone Dextrose (YPD) broth, and agar were purchased from BD Difco.

LB broth, MRS broth, and YPD broth were prepared with dH_2_O. All media-agar plates (LB-agar, MRS-agar, YPD-agar) were prepared with dH_2_O and 1.5 wt% agar. All growth media and PBS were sterilized by autoclaving at 120 °C for 20 min and then stored at room temperature.

### Cell culture

#### Bacterial culture

ABU 83972 strain of *E. coli* was used for Gram-negative bacteria release studies. Bacterial cultures were grown in LB broth. Initially, *E. coli* from glycerol stock was streaked onto LB-agar plates and incubated at 37 °C for 24 h. A single colony from the LB-agar plate was added to 25 mL of LB broth and incubated at 37 °C at 200 rpm (n = 3). After 24 h of incubation, the cultures were centrifuged at 4000 rpm for 10 min at room temperature, and the supernatants were removed. The bacterial cultures were then washed thrice with 25 mL of PBS to remove the culture media. The bacterial suspension was then diluted and adjusted to an optical density of 3.0 at 600 nm using a UV-vis spectrophotometer (Genesys 40, Thermo Scientific). To confirm the viable cell count, the solution was further diluted and plated on LB-agar plates. These plates were then incubated at 37 °C for 24 h and the CFUs were counted. The bacterial suspensions had ∼ 2 × 10^5^ CFU μL^−1^, which was then further diluted or concentrated for preparing ELMs.

For microscopy, ABU 83972 which expresses red fluorescence protein (RFP) was used. All the culture and growth were performed using the same procedure mentioned above. Except, LB-amp broth and LB-amp agar plates (50 μg of ampicillin per mL of media) were used instead of LB broth and LB-agar plates. *L. paracasei* was used for Gram-positive bacterial release studies. Initially, *L. paracasei* from glycerol stock was streaked onto MRS-agar plates. These plates were then incubated at 37 °C with 5% CO_2_ for 48 h. A single colony from the MRS-agar plate was added to 25 mL of MRS broth and incubated at 37 °C with 5% CO_2_ (*n* = 3). After 24 h of incubation, the cultures were centrifuged at 4000 rpm for 10 min at room temperature, and the supernatants were removed. The cells were then washed thrice with PBS. The bacterial suspension was then diluted and adjusted the optical density at 600 nm to 1.7 using a UV-vis spectrophotometer. The solution was then further diluted and plated on MRS-agar plates. These plates were then incubated at 37 °C and 5% CO_2_ for 48 h and the CFUs were counted. The bacterial suspensions had ∼ 1 × 10^5^ cells μL^−1^, which was then further diluted or concentrated for preparing ELMs.

#### Yeast culture

*S. cerevisiae* was used for yeast release studies. Initially, 10 mg of dry *S. cerevisiae* (Fleischmann’s Yeast) were dissolved in 1 mL of sterile dH_2_O, and the solution was streaked onto YPD-agar plates. These plates were then incubated at 30 °C for 48 h. A single colony from the YPD-agar plate was added to 25 mL of YPD broth and incubated at 30 °C in aerobic conditions with constant shaking at 200 rpm (n = 3). After 24 h of incubation, the cultures were centrifuged at 4000 rpm for 10 min at room temperature, and the supernatants were removed. The cells were then washed thrice with PBS. The yeast suspension was then diluted and the optical density at 600 nm was measured using a UV-vis spectrophotometer. For all replicates, the dilutions were adjusted to obtain an optical density of 1.4 at 600 nm. The solution was then further diluted and plated on YPD-agar plates. These plates were then incubated at 30 °C for 48 h and the CFUs were counted. The yeast suspensions had ∼ 1 × 10^5^ cells μL^−1^, which was then further diluted or concentrated for preparing ELMs.

### Preparation of ELMs

#### ELMs loaded with *E. coli*

he ELMs were prepared by free radical polymerization of HEA (monomer) and BIS (crosslinker). LAP was used as a photoinitiator for this polymerization. Stock solutions of BIS (0.02 g mL^−1^ and 0.04 g mL^−1^) and LAP (0.02 g mL^−1^) were prepared in dH_2_O. All chemicals and stock solutions were filter sterilized (polyethersulfone, 0.2 μm) before ELM preparation. ELMs with varying stiffnesses were prepared with a cell loading of 1 × 10^4^ cells μL^−1^. The ELMs with low-stiffness hydrogel (10 HEA / 0.1 BIS), medium-stiffness hydrogel (15 HEA / 0.5 BIS), and stiff hydrogel (20 HEA / 2.0 BIS) were prepared using 10 wt% HEA and 0.1 wt% BIS; 15 wt% HEA and 0.5 wt% BIS; 20 wt% HEA and 2.0 wt% BIS, respectively. The concentration of each component denotes the percentage by weight in the entire pre-gel solution, including cells. All ELMs were prepared with 0.04 wt% LAP. The remaining fraction of the pre-gel solution was filled with an equivalent mass of LB media. ELMs with varying cell loadings were prepared with medium stiffness hydrogels (15 HEA / 0.5 BIS). of 1 × 10^4^ cells μL^−1^. The ELMs with low, medium, and high cell loading were prepared with 1 × 10^0^ cells μL^−1^, 1 × 10^4^ cells μL^−1^, and 1 × 10^8^ cells μL^−1^, respectively. In these formulations, as the biomass was increased, the LB media volume was reduced, such that the total pre-gel solution amounted to 100 wt%. All the solutions were then vortexed for ∼5 s. After preparation, these pre-gel solutions were filled into a polyethylene tubing (PE50, Braintree Scientific) (Method #1) and exposed to UV irradiation (UVP Crosslinker CL-3000, Analytik Jena) of 365 nm at an intensity of 1.2 mW cm^−2^ for 2 min to polymerize. The polymerized ELMs were then dispensed from the tubing with a sterile 27G needle, trimmed to 4 mm length with razor blades, and washed thrice with PBS to remove unpolymerized monomer residues. The volume of the resulting ELMs was ∼1 μL.

For preparing ELMs with different volumes or different surface area (S.A.) to volume (V) ratios (S.A./V), medium cell loading (1 × 10^4^ cells μL^−1^) and medium stiffness hydrogel formulation (15 HEA / 0.5 BIS) were used. These ELMs were prepared by polymerizing the pre-gel solutions in a mold (Method #2). Molds were prepared by using two glass slides (75 mm × 50 mm × 1 mm) treated with a water-repellent spray (Rain-X Original), separated by a spacer. The thickness of the ELMs was defined by the thickness of the spacers. Different thicknesses of spacers that were used include 0.125 mm, 0.2 mm, 0.25 mm, 0.5 mm, 1 mm, 1.4 mm, and 2 mm. Upon the preparation of pre-gel solutions, the solutions were filled into molds and exposed to UV irradiation of 365 nm at an intensity of 1.2 mW cm^−2^ for 2 min to polymerize. The molds were flipped every 30 s during polymerization. After polymerization, the samples were removed from the glass slides and cut to the required diameter using biopsy punches and washed thrice with PBS to remove unpolymerized monomer residues. Different diameters of biopsy punches used were 2.5 mm, 4 mm, 5 mm, and 12 mm.

For preparing ELMs with single-colony, a cell loading of 1 × 10^0^ cells μL^−1^ was used to prepare the pre-gel solutions. The ELMs were prepared using molds by following the same procedure (Method #2). The prepared ELMs were incubated at 37 °C in aerobic conditions with constant shaking at 200 rpm for 24 h. The 1 day-grown ELMs with no colony, >1 colony, or any colonies near the surface of the hydrogels were not used for further experimentation. For growth, all ELMs were incubated in 10 mL LB-amp media and the media was refreshed every 24 h for 10 days.

PEGDA ELMs were prepared by free radical polymerization of PEGDA (monomer) in aqueous solution. LAP was used as a photoinitiator for this polymerization. For preparing the pre-gel solution, a medium cell loading (1 × 10^4^ cells μL^−1^) of *E. coli* and 10 wt% PEGDA (monomer) were used. The pre-gel solutions were then polymerized using the same procedure (Method #1).

Cell-free hydrogel samples used for mechanical characterization were also prepared using this approach (Method #2), but no cells were added to the pre-gel solution.

### *L. paracasei* loaded ELMs and *S. cerevisiae* loaded ELMs

The ELMs were prepared using the same procedure (Method #1), except ELMs with varying stiffnesses were prepared with a cell loading of 1 × 10^3^ cells μL^−1^; and ELMs with varying cell loading were prepared with a cell loading of 1 × 10^0^ cells μL^−1^, 1 × 10^3^ cells μL^−1^, and 1 × 10^6^ cells μL^−1^. During preparation, MRS media was used for *L. paracasei-*loaded ELMs, and YPD media was used for *S. cerevisiae*-loaded ELMs.

### Imaging and quantification of *E. coli* colony growth within ELMs

ELMs loaded with RFP expressing ABU 83972 (*E. coli*) were imaged using a confocal microscope (SP8, Leica Microsystems). The ELMs were exposed to a laser wavelength of 554 nm and power of 2%, and the detection wavelength was 540 – 650 nm, and shown with a green pseudo-color. Images were acquired with a resolution of 512 × 512 pixels using a 10x lens. Z-stacks of 500 μm were taken in steps of 0.2 μm.

The z-stack images were processed using Imaris software (Version 10.1, Oxford Instruments). The z-stack images were converted into 3D surfaces using the Imaris surface tool. The surfaces were generated with a smooth function set to 2 μm, background threshold to 10 μm, and the minimum surface voxel limit to 10. The volume of all bacterial colonies within the field of view was calculated.

### Quantification of bacteria/yeast release from ELMs

To quantify the release of *E. coli* from ELMs, the samples were placed in a 50 mL centrifuge tube with 10 mL of LB media and incubated at 37 °C in aerobic conditions with constant shaking at 200 rpm for 24 h. After 2 h of incubation, an aliquot of the media was diluted (in 10-fold dilutions, as necessary) and plated on LB-agar plates using an automatic plater (easySpiral, Interscience). These plates were then incubated at 37 °C overnight and the CFUs were counted using an automatic colony counter (Scan 300, Interscience). Every 24 h, the ELMs were removed from the LB media and washed thrice with PBS (each wash comprised vortexing the ELM with 10 mL of PBS at a medium speed for 20–30 s using a vortex mixer). The ELMs were then transferred to 10 mL of fresh LB media and incubated in the same conditions. After 2 h of incubation, the cell release was measured using the same procedure. The release of *E. coli* from the ELMs was measured for 10 days.

To quantify the release of *L. paracasei* from ELMs, the samples were placed in 10 mL of MRS media and incubated at 37 °C with 5% CO_2_ and no shaking for 24 h. After 6 h of incubation, an aliquot of the media was diluted and plated on MRS-agar plates using the automatic plater. These plates were then incubated at 37 °C with 5% CO_2_ for 48 h and the CFUs were counted using the automatic colony counter. Every 24 h, the ELMs were removed from the media, washed thrice with PBS, transferred to a fresh media, and incubated in the same conditions. After 6 h of incubation, the cell release was measured using the same procedure. The release of *L. paracasei* from the ELMs was measured for 10 days.

To quantify the release of *S. cerevisiae* from ELMs, the samples were placed in 10 mL of YPD media and incubated at 30 °C in aerobic conditions with constant shaking at 200 rpm for 24 h. After 6 h of incubation, an aliquot of the media was diluted and plated on YPD-agar plates using the Copacabana method. These plates were then incubated at 30 °C for 48 h and the CFUs were counted manually. Every 24 h, the ELMs were removed from the media, washed thrice with PBS, transferred to a fresh media, and incubated in the same conditions. After 6 h of incubation, the cell release was measured using the same procedure. The release of *S. cerevisiae* from the ELMs was measured for 10 days.

### Mechanical characterization

Cell-free hydrogel samples (12 mm in diameter, 2 mm in thickness) were equilibrated in LB media at 4 °C for 24 h. A uniaxial compression test was performed at room temperature, using a dynamic mechanical analyzer (RSA-G2, TA Instruments). The samples were loaded between the plates and ∼30 mL of LB media was added to the immersion system. The compression was then performed at a strain rate of 0.05 mm s^−1^ under immersion conditions. Strains between 0.2% and 5% were used to calculate Young’s modulus, as the stress-strain response in the region was linear.

### Monomer/crosslinker toxicity

The toxicity of the monomer and crosslinker was determined for *E. coli*, *L. paracasei*, and *S. cerevisiae*. To quantify the CFUs after exposure of each cell loading to each monomer/crosslinker concentration, all ELM formulations were prepared as described in the preparation of the ELM section, except that LAP was omitted to prevent gelation. Control solutions were prepared by replacing all monomers with LB media. After 2 min of exposure, an aliquot of the formulation was diluted (as necessary) and plated on LB-agar plates. These plates were incubated at 37 °C overnight and CFUs were counted.

To determine the toxicity of monomer and crosslinker for *L. paracasei*, the same procedure was followed using MRS media and MRS-agar plates instead of LB media and LB-agar plates, and the plates were incubated at 37 °C with 5% CO_2_ for 48 h before counting CFUs. Similarly, for *S. cerevisiae*, the same procedure was followed using YPD media and YPD-agar plates, and the plates were incubated at 30 °C for 48 h before counting CFUs.

### Quantification of bacteria present within ELMs

ELMs were washed thrice with PBS and placed in 3 mL of PBS for 30 min. ELMs were then homogenized using a handheld homogenizer blade (Homogenizer 150, Fisherbrand) at high speed for ∼30 s (until no visible hydrogel fragments remained). The homogenizer blade was decontaminated with 70% EtOH and washed with sterile dH_2_O between samples. After homogenization, an aliquot of the solution was diluted (as necessary) and plated on LB-agar plates. These plates were then incubated at 37 °C overnight and the CFUs were counted.

### Quantification of bacterial proliferation from released cells

5 day-grown ELMs were washed thrice with PBS and incubated in 10 mL of fresh LB media at 37 °C and 200 rpm. In condition 1, the ELMs were removed after 30 min and the remaining media was incubated in the same conditions for the next 90 min. In condition 2, the ELMs remained in the media for the entire 2 h. For both conditions, aliquots of the media at both 30 min and 120 min were diluted (as necessary) and plated on LB-agar plates. These plates were then incubated at 37 °C overnight and the CFUs were counted.

### Statistical analysis

Statistical analysis was performed using GraphPad Prism (Version 10.2.3). Data are shown as the mean ± standard deviation. For all statistical tests, *P* < 0.05 was set for statistical significance. Single comparisons were performed using a two-tailed Student’s *t*-test (unpaired), and multiple comparisons were performed using a one-way analysis of variance (ANOVA) with a post-hoc Tukey test.

## ACKNOWLEDGEMENTS

Research reported in this publication was partially supported by the National Institute of Biomedical Imaging and Bioengineering of the National Institutes of Health under Award Number R56EB032395 (T.H.W., S.S., and P.E.Z). This material is also partially based upon work supported by the National Institutes of Health under Grant No. DK114224 (S.S). The Funders had no role in study design, data collection and analysis, decision to publish, or preparation of the manuscript. The content is solely the responsibility of the authors and does not necessarily represent the official views of the funders. Some of the graphics were created with BioRender.com.

## Notes

### Competing Interest Statement

The authors have declared no competing interest.

## REFERENCES

1. Yoshida, R., Sakai, K., Okano, T. & Sakurai, Y. A New Model for Zero-Order Drug Release I. Hydrophobic Drug Release from Hydrophilic Polymeric Matrices. Polym. J. 23, 1111–1121 (1991).

2. Vargason, A. M., Anselmo, A. C. & Mitragotri, S. The evolution of commercial drug delivery technologies. *Nat*. Biomed. Eng. 5, 951–967 (2021).

3. Cook, M. T., Tzortzis, G., Charalampopoulos, D. & Khutoryanskiy, V. V. Microencapsulation of probiotics for gastrointestinal delivery. J. Control. Release 162, 56–67 (2012).

4. Sharma, H., Sharma, S., Bajwa, J., Chugh, R. & Kumar, D. Polymeric carriers in probiotic delivery system. Carbohydr. Polym. Technol. Appl. 5, 100301 (2023).

5. Stamatopoulos, K., Kafourou, V., Batchelor, H. K. & Konteles, S. J. Sporopollenin Exine Microcapsules as Potential Intestinal Delivery System of Probiotics. Small 17, e2004573 (2021).

6. Laracuente, M.-L., Yu, M. H. & McHugh, K. J. Zero-order drug delivery: State of the art and future prospects. J. Control. Release 327, 834–856 (2020).

7. Narasimhan, B. & Langer, R. Zero-order release of micro- and macromolecules from polymeric devices: the role of the burst effect. J. Control. Release 47, 13–20 (1997).

8. Razavi, S., Janfaza, S., Tasnim, N., Gibson, D. L. & Hoorfar, M. Microencapsulating polymers for probiotics delivery systems: Preparation, characterization, and applications. Food Hydrocoll. 120, 106882 (2021).

9. Roos, V., Ulett, G. C., Schembri, M. A. & Klemm, P. The Asymptomatic Bacteriuria Escherichia coli Strain 83972 Outcompetes Uropathogenic E. coli Strains in Human Urine. Infect Immun 74, 615–624 (2006).

10. Falagas, M. E., Betsi, G. I., Tokas, T. & Athanasiou, S. Probiotics for Prevention of Recurrent Urinary Tract Infections in Women: A Review of the Evidence from Microbiological and Clinical Studies. Drugs 66, 1253–1261 (2006).

11. Wang, F. et al. Probiotics in Helicobacter pylori eradication therapy: Systematic review and network meta-analysis. Clin. Res. Hepatol. Gastroenterol. 41, 466–475 (2017).

12. Song, C. H. et al. Lactobacillus crispatus Limits Bladder Uropathogenic E. coli Infection by Triggering a Host Type I Interferon Response. Proc. Natl. Acad. Sci. United States Am. 119, e2117904119 (2022).

13. Duraj-Thatte, A. M. et al. Modulating bacterial and gut mucosal interactions with engineered biofilm matrix proteins. Sci. Rep. 8, 3475 (2018).

14. Lehtoranta, L., Ala-Jaakkola, R., Laitila, A. & Maukonen, J. Healthy Vaginal Microbiota and Influence of Probiotics Across the Female Life Span. Front. Microbiol. 13, 819958 (2022).

15. Shen, N. T. et al. Timely Use of Probiotics in Hospitalized Adults Prevents Clostridium difficile Infection: A Systematic Review With Meta-Regression Analysis. Gastroenterology 152, 1889–1900.e9 (2017).

16. George, I., Kalairaj, M. S., Zimmern, P. E., Ware, T. H. & Subashchandrabose, S. Competitive fitness of asymptomatic bacteriuria E. coli strain 83972 against uropathogens in human urine. Infect. Immun. e0017324 (2024) doi:10.1128/iai.00173-24.

17. Miller, L. E., Ouwehand, A. C. & Ibarra, A. Effects of probiotic-containing products on stool frequency and intestinal transit in constipated adults: systematic review and meta-analysis of randomized controlled trials. Ann. Gastroenterol. 30, 629–639 (2017).

18. Akbari, V. & Hendijani, F. Effects of probiotic supplementation in patients with type 2 diabetes: systematic review and meta-analysis. Nutr. Rev. 74, 774–784 (2016).

19. Saxelin, M., Tynkkynen, S., Mattila-Sandholm, T. & Vos, W. M. de. Probiotic and other functional microbes: from markets to mechanisms. Curr. Opin. Biotechnol. 16, 204–211 (2005).

20. Azevedo, M. S., Sande, F. H. van de, Romano, A. R. & Cenci, M. S. Microcosm Biofilms Originating from Children with Different Caries Experience Have Similar Cariogenicity under Successive Sucrose Challenges. Caries Res. 45, 510–517 (2011).

21. Lerner, A., Shoenfeld, Y. & Matthias, T. Probiotics: If It Does Not Help It Does Not Do Any Harm. Really? Microorganisms 7, 104 (2019).

22. Suez, J., Zmora, N., Segal, E. & Elinav, E. The pros, cons, and many unknowns of probiotics. Nat. Med. 25, 716–729 (2019).

23. Jacobsen, C., García-Moreno, P. J., Mendes, A. C., Mateiu, R. V. & Chronakis, I. S. Use of Electrohydrodynamic Processing for Encapsulation of Sensitive Bioactive Compounds and Applications in Food. Annu. Rev. Food Sci. Technol. 9, 1–25 (2017).

24. Diep, E. & Schiffman, J. D. Encapsulating bacteria in alginate-based electrospun nanofibers. Biomater. Sci. 9, 4364–4373 (2021).

25. Lau, L. Y. J. & Quek, S. Y. Probiotics: Health benefits, food application, and colonization in the human gastrointestinal tract. Food Bioeng. 3, 41–64 (2024).

26. Gu, Q. et al. Encapsulation of multiple probiotics, synbiotics, or nutrabiotics for improved health effects: A review. Adv. Colloid Interface Sci. 309, 102781 (2022).

27. Han, S. et al. Probiotic Gastrointestinal Transit and Colonization After Oral Administration: A Long Journey. Front. Cell. Infect. Microbiol. 11, 609722 (2021).

28. García-Cayuela, T. et al. Adhesion abilities of dairy Lactobacillus plantarum strains showing an aggregation phenotype. Food Res. Int. 57, 44–50 (2014).

29. Cao, Z., Wang, X., Pang, Y., Cheng, S. & Liu, J. Biointerfacial self-assembly generates lipid membrane coated bacteria for enhanced oral delivery and treatment. Nat. Commun. 10, 5783 (2019).

30. Anselmo, A. C., McHugh, K. J., Webster, J., Langer, R. & Jaklenec, A. Layer-by-Layer Encapsulation of Probiotics for Delivery to the Microbiome. Adv. Mater. 28, 9486–9490 (2016).

31. Praveschotinunt, P. et al. Engineered E. coli Nissle 1917 for the delivery of matrix-tethered therapeutic domains to the gut. Nat. Commun. 10, 5580 (2019).

32. Heavey, M. K. et al. Targeted delivery of the probiotic Saccharomyces boulardii to the extracellular matrix enhances gut residence time and recovery in murine colitis. Nat. Commun. 15, 3784 (2024).

33. Wang, X. et al. Bioinspired oral delivery of gut microbiota by self-coating with biofilms. Sci. Adv. 6, eabb1952 (2020).

34. O’Brien, C. E. et al. Early probiotic supplementation with B. infantis in breastfed infants leads to persistent colonization at 1 year. Pediatr. Res. 91, 627–636 (2022).

35. Stærk, K. et al. Bladder catheterization improves bacterial interference with asymptomatic Escherichia coli 83972 in an experimental porcine model of urinary tract infection. J. Infect. Dis. jiae404 (2024) doi:10.1093/infdis/jiae404.

36. Luan, Q. et al. Controlled Nutrient Delivery through a pH-Responsive Wood Vehicle. ACS Nano 16, 2198–2208 (2022).

37. Rajab, S., Tabandeh, F., Shahraky, M. K. & Alahyaribeik, S. The effect of lactobacillus cell size on its probiotic characteristics. Anaerobe 62, 102103 (2020).

38. Sankaran, S., Becker, J., Wittmann, C. & Campo, A. del. Optoregulated Drug Release from an Engineered Living Material: Self-Replenishing Drug Depots for Long-Term, Light-Regulated Delivery. Small 15, e1804717 (2019).

39. Rodrigo-Navarro, A., Sankaran, S., Dalby, M. J., Campo, A. del & Salmeron-Sanchez, M. Engineered living biomaterials. Nat. Rev. Mater. 6, 1175–1190 (2021).

40. Rivera-Tarazona, L. K., Bhat, V. D., Kim, H., Campbell, Z. T. & Ware, T. H. Shape-morphing living composites. Sci. Adv. 6, eaax8582 (2020).

41. Bhusari, S., Sankaran, S. & Campo, A. del. Regulating Bacterial Behavior within Hydrogels of Tunable Viscoelasticity. Adv. Sci. 9, 2106026 (2022).

42. Saha, A. et al. Additive Manufacturing of Catalytically Active Living Materials. ACS Appl. Mater. Interfaces 10, 13373–13380 (2018).

43. Molinari, S., Tesoriero, R. F. & Ajo-Franklin, C. M. Bottom-up approaches to engineered living materials: Challenges and future directions. Matter 4, 3095–3120 (2021).

44. Rivera-Tarazona, L. K. et al. Controlling shape morphing and cell release in engineered living materials. Biomater. Adv. 143, 213182 (2022).

45. Bober, J. R., Beisel, C. L. & Nair, N. U. Synthetic Biology Approaches to Engineer Probiotics and Members of the Human Microbiota for Biomedical Applications. Annu. Rev. Biomed. Eng. 20, 1–24 (2018).

46. Tuson, H. H. et al. Measuring the stiffness of bacterial cells from growth rates in hydrogels of tunable elasticity. Mol. Microbiol. 84, 874–891 (2012).

47. Tang, T.-C. et al. Hydrogel-based biocontainment of bacteria for continuous sensing and computation. Nat. Chem. Biol. 17, 724–731 (2021).

48. Lee, J. W., Chan, C. T. Y., Slomovic, S. & Collins, J. J. Next-generation biocontainment systems for engineered organisms. Nat. Chem. Biol. 14, 530–537 (2017).

49. Lacroix, C., Paquin, C. & Arnaud, J.-P. Batch fermentation with entrapped growing cells ofLactobacillus casei. Appl. Microbiol. Biotechnol. 32, 403–408 (1990).

50 Zhang, Q., et al. Morphogenesis and cell ordering in confined bacterial biofilms. Proc. Natl. Acad. Sci. 118, e2107107118 (2021).

51. Lee, J. et al. Determining the Young’s Modulus of the Bacterial Cell Envelope. ACS Biomater. Sci. Eng. 10, 2956–2966 (2024).

52. Kang, J., Wang, C. & Cai, S. Cavitation to fracture transition in a soft solid. Soft Matter 13, 6372–6376 (2017).

53. Kato, M., Fujii, T. & Onaka, S. Elastic strain energies of sphere, plate and needle inclusions. Mater. Sci. Eng.: A 211, 95–103 (1996).

54. Hill, C. et al. The International Scientific Association for Probiotics and Prebiotics consensus statement on the scope and appropriate use of the term probiotic. Nat. Rev. Gastroenterol. Hepatol. 11, 506–514 (2014).

55. Latif, A. et al. Probiotics: mechanism of action, health benefits and their application in food industries. Front. Microbiol. 14, 1216674 (2023).

56. Oh, J. et al. Growth, Distribution, and Photosynthesis of Chlamydomonas Reinhardtii in 3D Hydrogels. Adv. Mater. 36, e2305505 (2024).

57. Śliżewska, K. & Chlebicz-Wójcik, A. Growth Kinetics of Probiotic Lactobacillus Strains in the Alternative, Cost-Efficient Semi-Solid Fermentation Medium. Biology 9, 423 (2020).

58. Fukuda, N. Apparent diameter and cell density of yeast strains with different ploidy. Sci. Rep. 13, 1513 (2023).

59. Sundén, F., Håkansson, L., Ljunggren, E. & Wullt, B. Escherichia coli 83972 Bacteriuria Protects Against Recurrent Lower Urinary Tract Infections in Patients With Incomplete Bladder Emptying. J. Urol. 184, 179–185 (2010).

60. Chee, W. J. Y., Chew, S. Y. & Than, L. T. L. Vaginal microbiota and the potential of Lactobacillus derivatives in maintaining vaginal health. Microb. Cell Factories 19, 203 (2020).

61. Fakruddin, Md., Hossain, Md. N. & Ahmed, M. M. Antimicrobial and antioxidant activities of Saccharomyces cerevisiae IFST062013, a potential probiotic. BMC Complement. Altern. Med. 17, 64 (2017).

62. Gaziano, R., Sabbatini, S., Roselletti, E., Perito, S. & Monari, C. Saccharomyces cerevisiae-Based Probiotics as Novel Antimicrobial Agents to Prevent and Treat Vaginal Infections. Front. Microbiol. 11, 718 (2020).

63. Babaei, F., Mirzababaei, M., Mohammadi, G., Dargahi, L. & Nassiri-Asl, M. Saccharomyces boulardii attenuates lipopolysaccharide-induced anxiety-like behaviors in rats. Neurosci. Lett. 778, 136600 (2022).

64. Cook, C. M. et al. The probiotic Lacticaseibacillus paracasei strain Shirota (LcS) in a fermented milk beverage survives the gastrointestinal tract of generally healthy U.S. Adults. Int. J. Food Sci. Nutr. 74, 645–653 (2023).

65. Andersson, P. et al. Persistence of Escherichia coli bacteriuria is not determined by bacterial adherence. Infect. Immun. 59, 2915–2921 (1991).

66. Nagpal, R. et al. Human-origin probiotic cocktail increases short-chain fatty acid production via modulation of mice and human gut microbiome. Sci. Rep. 8, 12649 (2018).

