## Supplementary material for "Controlled release of microorganisms from engineered living materials": SI

**This document includes:**

Supporting Information Text

Supplementary Figures 1 to 12

**Supporting Information Text**

The cells released from the ELMs can proliferate in the growth media, but cell proliferation in the growth media does not significantly change the measured number of released cells in the first 2 h. For quantifying cell release, we measured the number of cells present in the media after 2 h of incubation. In this method, the viable cells released from the ELMs can continue to proliferate in the growth media. To better quantify the cells released as compared to cells proliferated in the media, we measured colony-forming units present in the media under two conditions (**Supplementary Fig. 7a**). Five day-grown ELMs were suspended in fresh LB media and incubated at 37 °C and 200 rpm. Then, in condition 1, the ELMs were removed after 30 min and the remaining media was incubated in the same conditions for the next 90 min. Here, the cells released from the ELMs in the first 30 min can further proliferate within the media for the next 90 min. The number of cells present in the media at 30 min did not increase significantly in the next 90 min (**Supplementary Fig. 7b**). On the other hand, in condition 2, the ELMs remained in the media for the entire 2 h, hence, the ELMs release cells for the entire 2 h. The number of cells present in the media at 30 min increased significantly in the next 90 min for all formulations ($P < 0.0001$ for low stiffness ELMs, $P < 0.01$ for medium stiffness ELMs, and $P < 0.001$ for high stiffness ELMs) (**Supplementary Fig. 7c**). These data collectively support that further proliferation of released cells does not significantly influence the number of cells measured at 2 h.

The proliferation of cells within ELMs is essential for cell release. We compared the cell release from 5 day-grown ELMs in shaking conditions (37 °C, 200 rpm) with static conditions (37 °C). There was no significant difference between the number of cells released by the ELMs in shaking and static conditions (**Supplementary Fig. 8**). This suggests that the cell release is driven by the forces associated with cell proliferation and not collisions with the container. Moreover, ELMs only release cells in growth media such as LB, and not in saline. However, when ELMs grown in LB are transferred to saline, lower cell release is observed which reduces dramatically with time. To better quantify this residual cell release, 5 day-grown ELMs were incubated in LB media and saline, and their cell release was compared at different time points (**Supplementary Fig. 9a**). During the first 30 min, there was no significant difference between the number of cells released in saline and LB media (**Supplementary Fig. 9b**). Additionally, in saline, the cells released in the first 30 min did not increase significantly during the next 90 min (**Supplementary Fig. 9c**). In contrast, in LB media, as discussed earlier, the cells released in the first 30 min significantly increased during the next 90 min. These data collectively suggest that the residual cell release plays a significant role in the first 30 min, whereas the forces associated with cell proliferation play a significant role during the next 90 min.

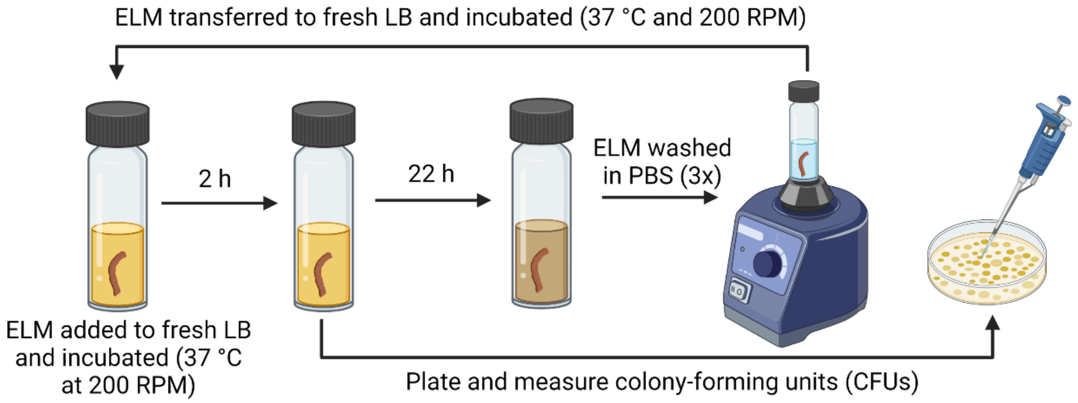


**Supplementary Fig. 1 |** Schematic showing the steps involved in quantifying cell (*E. coli*) release from ELMs.

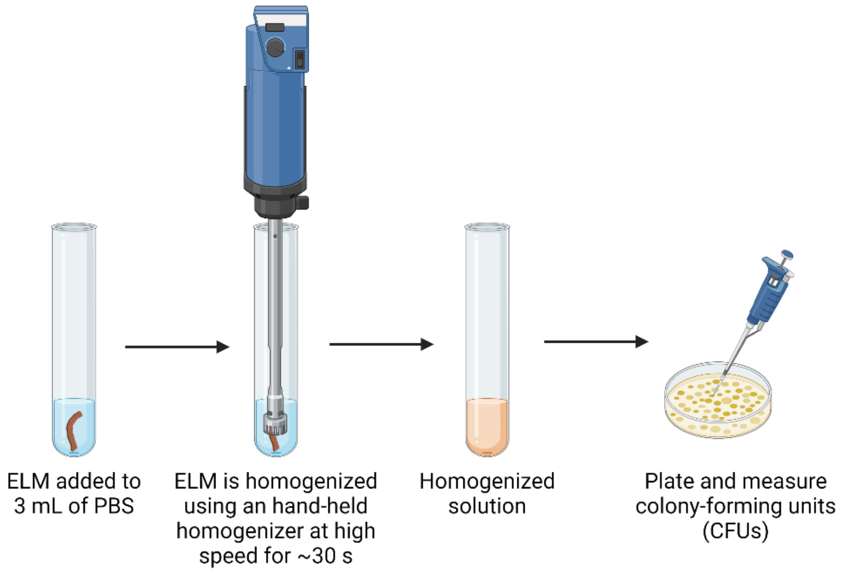


**Supplementary Fig. 2 |** Schematic showing the steps involved in quantifying cells (*E. coli*) present within ELMs.

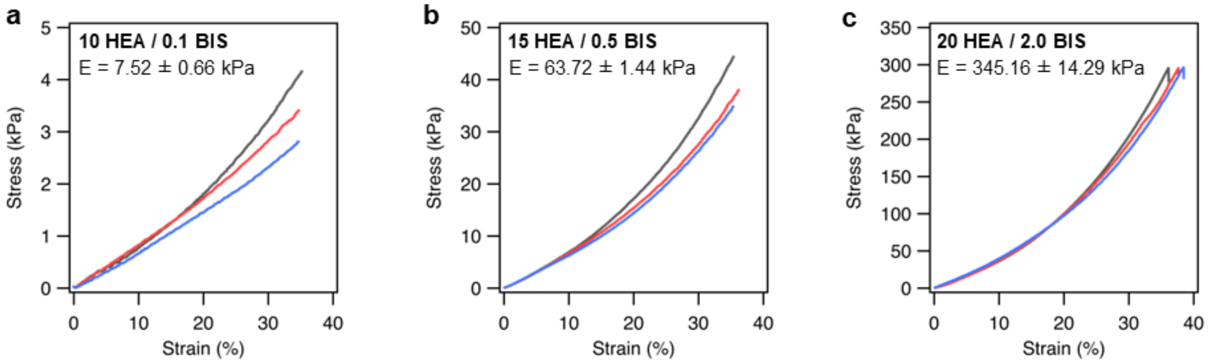


**Supplementary Fig. 3 |** Stress-strain curves of HEA / BIS hydrogels ($n$ = 3). **(a)** 10 HEA / 0.1 BIS, **(b)** 15 HEA/ 0.5 BIS, **(c)** 20 HEA / 2.0 BIS.

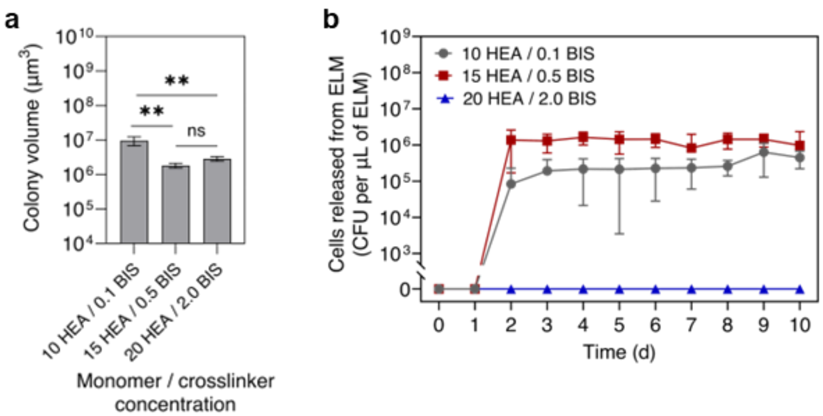


**Supplementary Fig. 4 |** Influence of hydrogel stiffness on fracture and cell release. **(a)** Colony volume as a function of different hydrogel stiffnesses (10 HEA / 0.1 BIS, 15 HEA/ 0.5 BIS, and 20 HEA / 2.0 BIS). **(b)** Cell release as a function of time from ELMs with different stiffnesses. All ELMs were 0.5 mm thick and loaded with a single *E. coli*. All data are presented as mean ± standard deviation ($n$ = 3). Statistical analysis was performed by a one-way ANOVA with post-hoc Tukey's test, ** $P < 0.01$, not significant (ns) for $P > 0.05$.

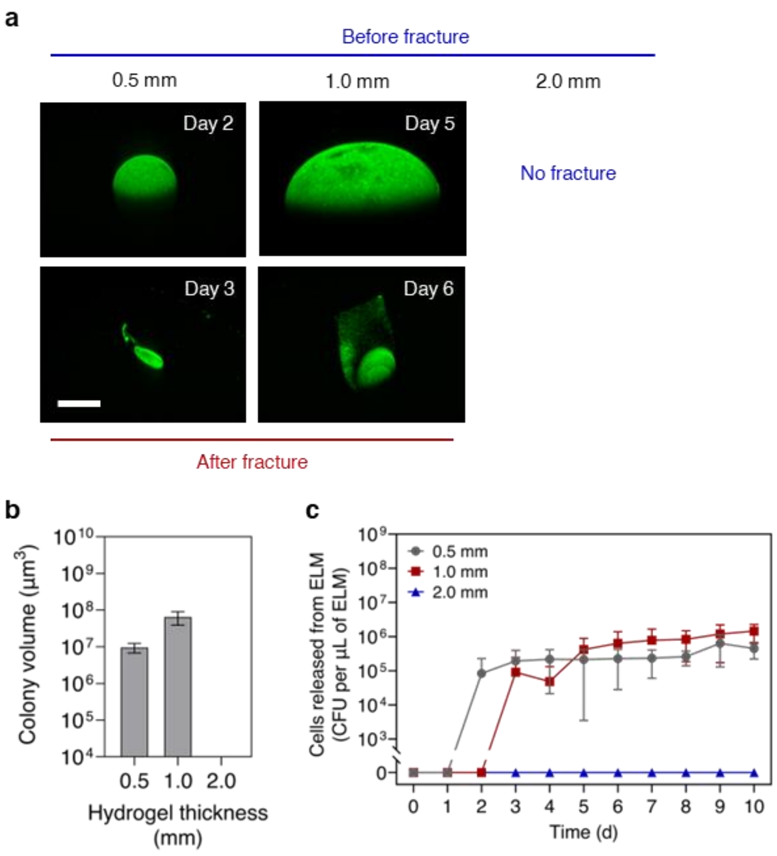


**Supplementary Fig. 5 |** Influence of hydrogel thickness on fracture and cell release. **(a)** Confocal microscopy z-stack images showing colony growth and fracture in ELMs with different thicknesses. Scale bar, 200 μm. **(b)** Colony volume as a function of different hydrogel thicknesses. **(c)** Cell release as a function of time from ELMs with different thicknesses (0.5, 1, and 2 mm). All ELMs were prepared with low-stiffness hydrogel (10 HEA / 0.1 BIS) and loaded with a single *E. coli*. All data are presented as mean ± standard deviation ($n = 3$).

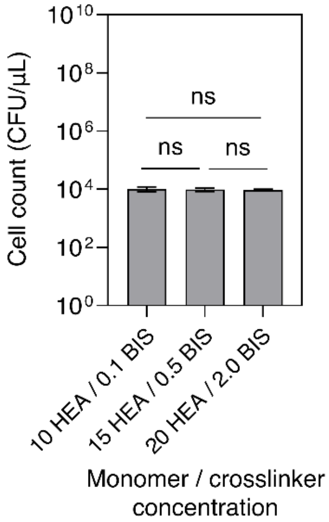


**Supplementary Fig. 6 |** *E. coli* viability as a function of exposure to formulations with different monomer and crosslinker content (10 HEA / 0.1 BIS, 15 HEA/ 0.5 BIS, and 20 HEA / 2.0 BIS). All data are presented as mean ± standard deviation ($n$ = 3). Statistical analysis was performed by a one-way ANOVA with post-hoc Tukey's test, not significant (ns) for $P > 0.05$.

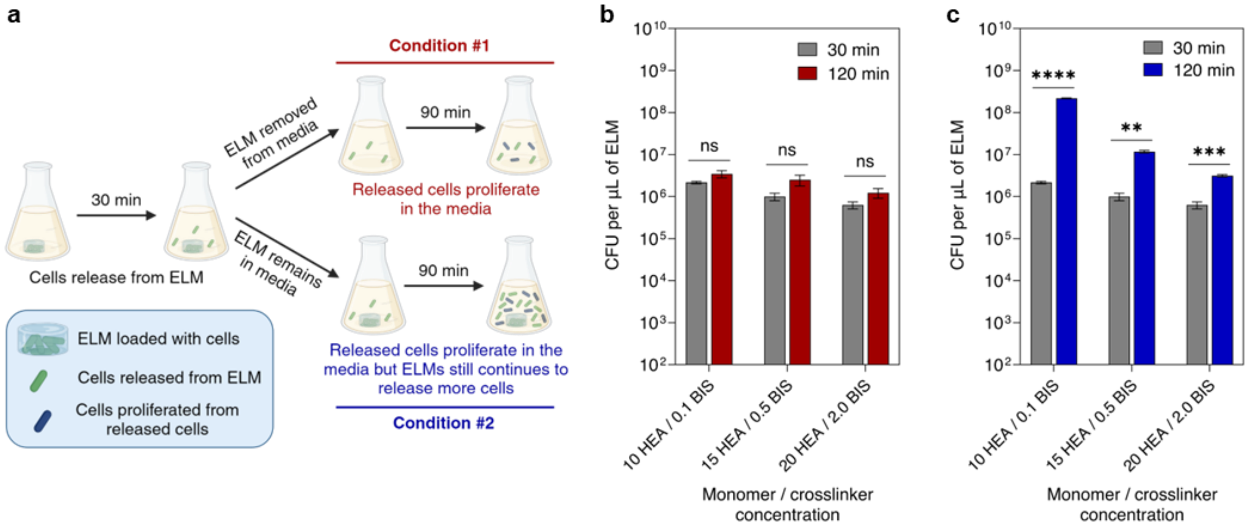


**Supplementary Fig. 7 |** Influence of cell proliferation from released cells. **(a)** Illustration of cell release and further proliferation of released cells (created with BioRender.com). Cells are released from ELMs in the first 30 min. In condition #1, ELMs are removed after 30 min: the released cells continue to proliferate during the next 90 min. In condition #2, ELMs remain in the media for 2 h: both released cells continue to proliferate, and ELMs continue to release cells for the next 90 min. **(b)** Cell count in condition #1. **(c)** Cell count in condition #2. All data are presented as mean ± standard deviation ($n$ = 3). Statistical analysis was performed by a two-tailed Student's $t$-test. Not significant (ns) for $P > 0.05$.

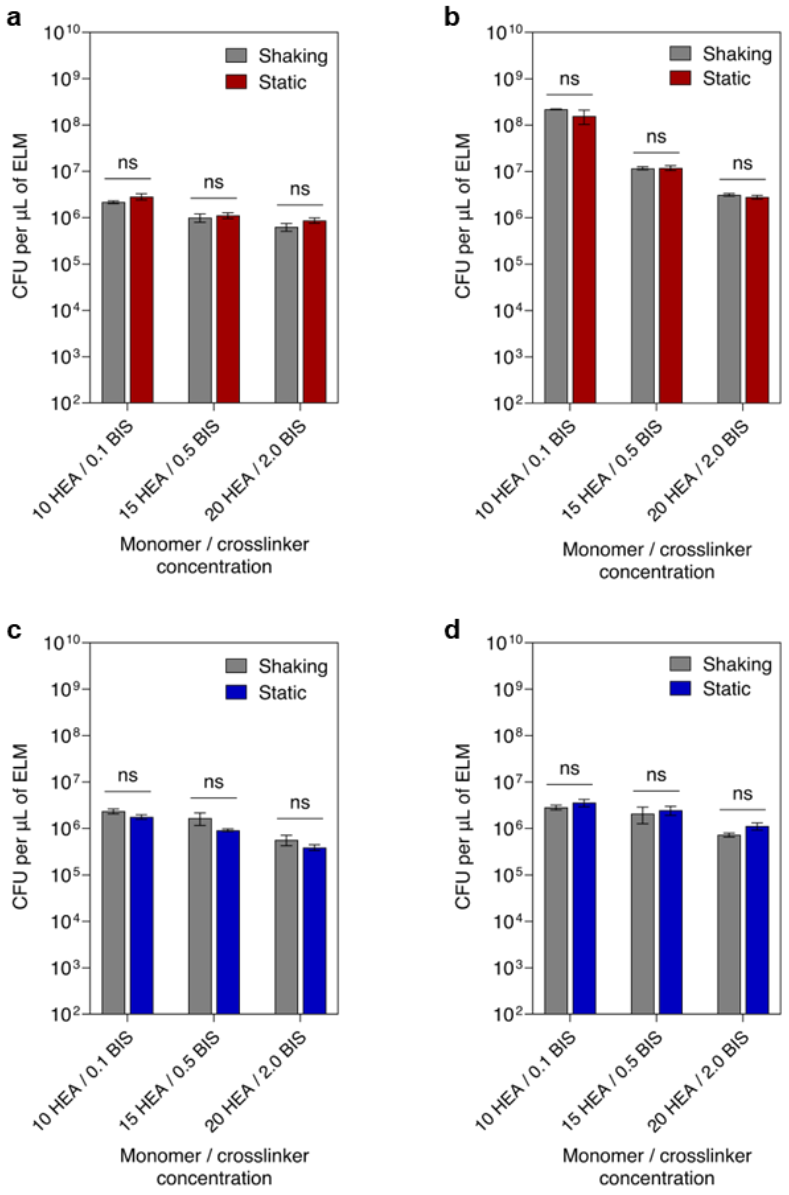


**Supplementary Fig. 8 |** Comparison of cell release from ELMs in shaking (37 °C, 200 rpm) vs static conditions (37 °C). **(a)** Cell release in LB for 30 min. **(b)** Cell release in LB for 2 h. **(c)** Cell release in PBS for 30 min. **(d)** Cell release in PBS for 2 h. ELMs were prepared with a cell loading of 1 × $10^4$ cells per µL of ELM and different hydrogel formulations (10 HEA / 0.1 BIS, 15 HEA / 0.5 BIS, 20 HEA / 2.0 BIS). All data are presented as mean ± standard deviation ($n$ = 3). Statistical analysis was performed by a two-tailed Student's $t$-test. Not significant (ns) for $P > 0.05$.

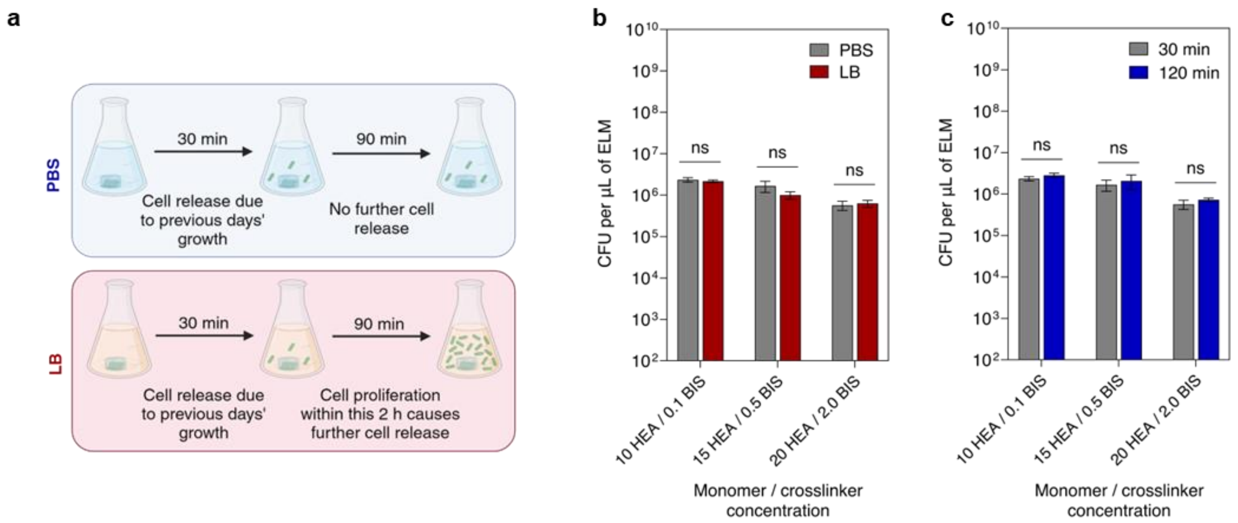


**Supplementary Fig. 9 | Comparison of cell release from ELMs in PBS and LB. (a)** Schematic showing the difference in cell release from ELMs when incubated in LB vs PBS (created with BioRender.com). **(b)** Comparison of cell release from ELMs in LB and PBS for 30 min. **(c)** Comparison of cell release in PBS for 30 min and 2 h. ELMs were prepared with a cell loading of $1 \times 10^4$ cells per µL of ELM and different hydrogel formulations (10 HEA / 0.1 BIS, 15 HEA / 0.5 BIS, 20 HEA / 2.0 BIS). All data are presented as mean ± standard deviation ($n$ = 3). Statistical analysis was performed by a two-tailed Student's *t*-test. Not significant (ns) for $P > 0.05$.

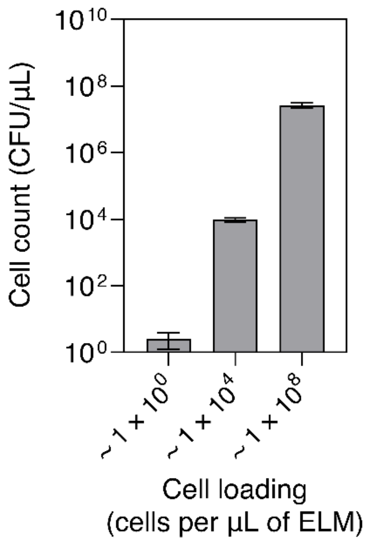


**Supplementary Fig. 10 |** *E. coli* viability as a function of different cell loading (1 × $10^0$ cells per µL of ELM, 1 × $10^4$ cells per µL of ELM, and 1 × $10^8$ cells per µL of ELM) when exposed to formulation (15 HEA/ 0.5 BIS). All data are presented as mean ± standard deviation ($n$ = 3).

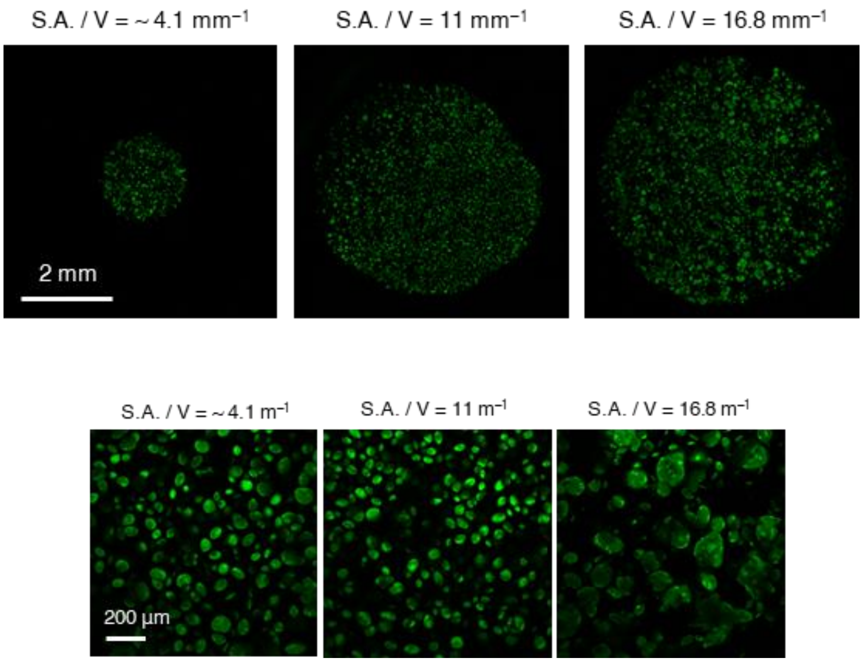


**Supplementary Fig. 11 |** Microscopy images showing the differences in colony morphologies as the S.A./ V ratios are varied (~4.1, 11, and 16.8 $mm^{-1}$). All ELMs were prepared with medium stiffness hydrogel (15 HEA / 0.5 BIS) and a cell loading of $1 \times 10^4$ cells per µL of ELM.

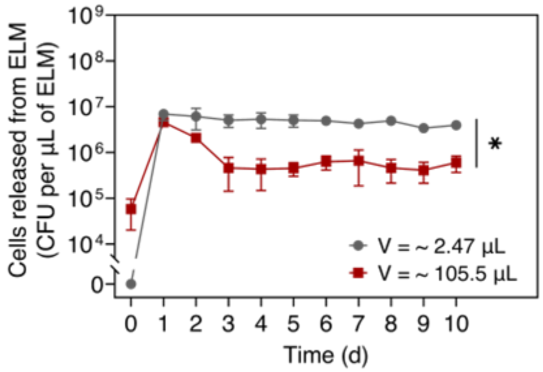


**Supplementary Fig. 12 |** Cell release as a function of time from ELMs of different sizes (V = ~2.47 and ~105.5 µL). All ELMs were prepared with medium stiffness hydrogel (15 HEA / 0.5 BIS) with a cell loading of $1 \times 10^4$ cells per µL of ELM and had a S.A./V ratio of ~4.1 $mm^{-1}$. All data are presented as mean ± standard deviation ($n$ = 3). Statistical analysis was performed by a two-tailed Student's $t$-test, * $P < 0.05$.
